# Minute-timescale simulations of G Protein Coupled Receptor A2A activation mechanism reveal a receptor pseudo-active state

**DOI:** 10.1101/2023.09.14.557711

**Authors:** Vincenzo Maria D’Amore, Paolo Conflitti, Luciana Marinelli, Vittorio Limongelli

## Abstract

G protein coupled receptors (GPCRs) are membrane proteins of greatest pharmacological relevance, targeted by over one third of marketed drugs. These receptors are activated by orthosteric ligands and undergo large conformational changes that lead to coupling diverse effector proteins. To achieve a fine regulation of the drug pharmacological response, it is imperative to shed light on the yet poorly understood aspects of GPCRs activation. In this work, we elucidate the entire activation mechanism of the adenosine A2A receptor (A2AR), a class A GPCR, performing minute timescale molecular dynamics and free energy calculations. We have explored the entire conformational landscape of A2AR in its basal apo form and in differently ligated conditions, elucidating the ligand intrinsic activity and the receptor’s lowest energy functional states. Among these is a novel pseudo-active state (pAs) of the A2AR apo form stabilised by specific “microswitch” residues interactions, including the salt bridge between the class A conserved residues R^5.66^ and E^6.30^. In the pAs state, A2AR is able to couple β-arrestin 1 over G proteins, providing unprecedented structural basis for receptor desensitization and G protein-alternative cellular pathways. Our simulation protocol is generalisable and can be applied to study the activation of any GPCR, resulting a precious tool for drug design and biased signaling studies.

## Introduction

G protein coupled receptors (GPCRs) are prominent pharmacological targets, representing 4% of the protein-coding genome and being targeted by almost 40% of the currently marketed drugs.^1,2^ In response to extracellular stimuli like hormones, neurotransmitters, and odorants, GPCRs regulate a plethora of biological functions, including vision, inflammation, and sensory perception.^1^ They show a conserved structural architecture arranged in seven transmembrane (TM) helices, connected through 3 extracellular (ECL) and 3 intracellular (ICL) loops.^3^ The GPCR barrel-like tertiary structure can be depicted in three main sections (Fig. 1): i) the orthosteric ligand binding site (OBS) at the extracellular region; ii) the connector; and iii) the intracellular binding site (IBS) where binding of effector proteins (aka transducers) occurr.^4^ The operating system is a ternary complex where GPCR is bound to a ligand and at the same time to a transducer that triggers the signal cascade inside the cell.^5^ The GPCRs are endowed with intrinsic functional dynamics and upon agonist binding the receptor undergoes large-scale conformational changes passing from the inactive to the active state. For instance, in Rhodopsin-family class A GPCRs, the two terminal states - i.e., active and inactive - differ for an outward/inward motion of the cytoplasmic end of TM6 (∼ 12-14 Å), accompanied by a rotation of the same helix around its axis (∼ 40°-50°) and slight shifts of the C-ter side of TM5 and TM7 (Fig. 1).^5–7^ In doing so, the GPCR IBS opens up, promoting the interaction with the transducer. On the other hand, minor differences are found with respect to the OBS by comparing active and inactive structures (in the range of 1.5-2 Å).^8,9^ Nonetheless, the receptor (de)activation appears to be regulated by a fine-grained allosteric communication between the two regions. Indeed, the binding of an agonist to OBS promotes the recruitment of an intracellular transducer at IBS; at the same time, the binary transducer-GPCR complex has higher affinity for agonist than the sole receptor.^10–12^ In addition, GPCRs can trigger a wide range of cellular pathways^10,13,14,15^ by coupling with diverse effector proteins such as G proteins (GPs)^13,15^, G protein-coupled receptor kinases (GRKs)^16^, proto-oncogene c-Src^17^ and arrestins^18,19^. In this scenario, a ligand might induce specific receptor conformations competent for the binding to a certain effector, thus selectively activating downstream cell signaling.^20–25^ This phenomenon, known as “biased signalling”^20,22^, brought to the fore new important implications for the pharmaceutical and clinical application of GPCR-targeting drugs.^20,23,26^ For instance, developing molecules capable of selectively activating or inhibiting a specific signalling cascade can yield a more targeted modulation of cell function with consequently reduced adverse effects,^27,28^ as recently reported for antidepressant drugs targeting the serotonin 5-HT_2A_ receptor,^29^ and analgesics targeting adenosine A1^30^ and μ opioid receptor^31^. Nevertheless, the rational design of “biased” ligands is hampered by the lack of structural information on all the functional conformations assumed by the GPCRs along their activation process. In fact, although the so-called “Resolution Revolution”^32^ in structural biology is constantly advancing the molecular understanding of the end states, other fundamental aspects of the GPCRs’ functional mechanism remain elusive, including the activation dynamics and the possible presence of receptor metastable, intermediate states. Atomistic simulations based on molecular dynamics (MD) techniques have demonstrated ability in detecting dynamic properties of the receptor, including the interaction with ligands and effectors.^9,31,33–50^. However, the long - ms - timescale and the complex allosteric network ruling GPCR activation make it elusive even to sophisticated simulation protocols.

**Figure 1.**
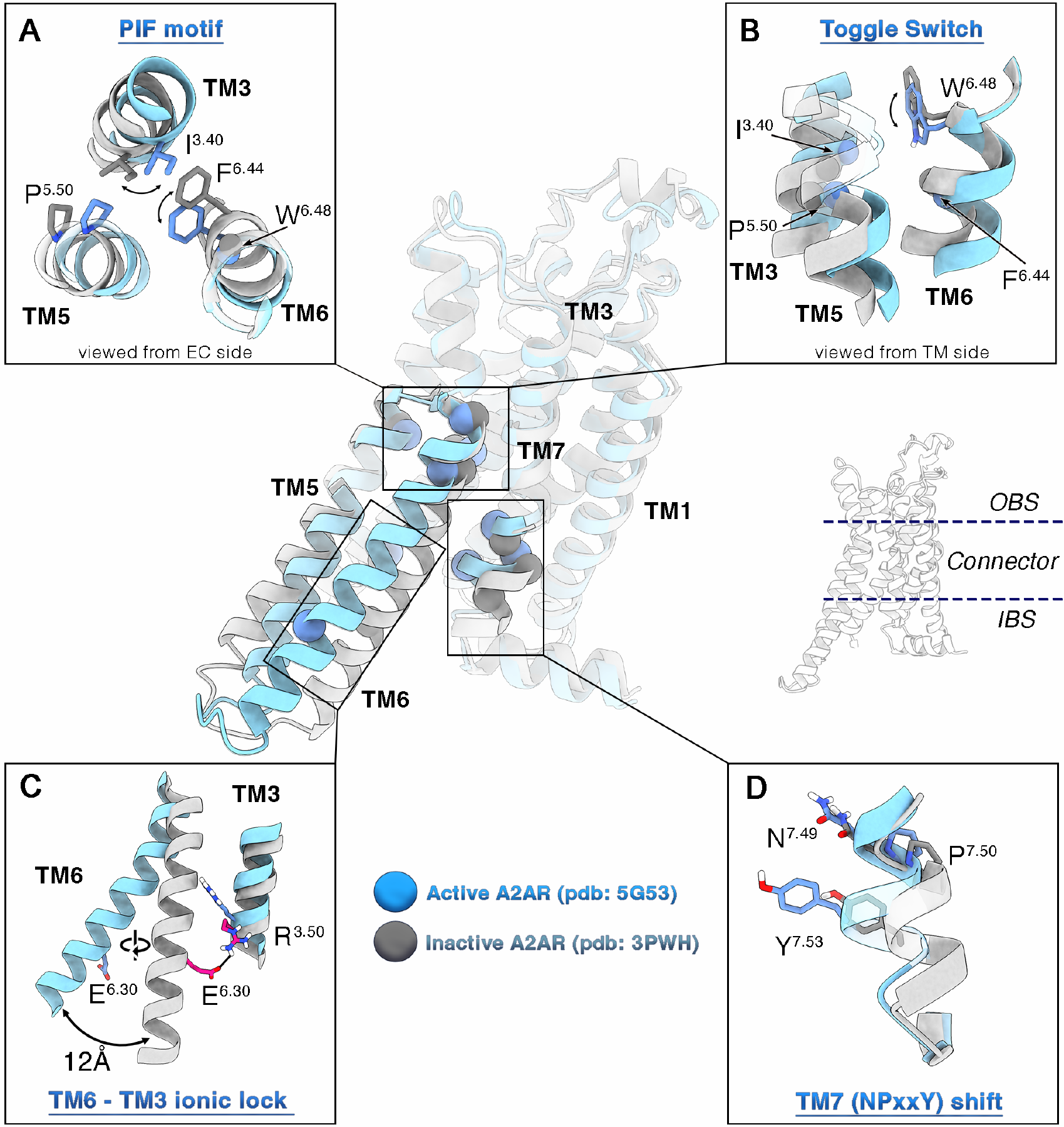
Main conformational changes occurring during GPCR activation. Active and inactive conformations of the A2AR are shown as cyan and silver cartoon, respectively. The four fundamental class A microswitches are highlighted as insets: the PIF motif (connector region, A), the W6.48 rotameric state “toggle switch” (OBS/connector region, B), the TM6-TM3 inactivating ionic lock (“IIL”, IBS region, C) and the NPxxY inward/outward shift (IBS region, D). The residues mainly involved in these transitions are shown as spheres (representing the a-carbon) onto the central representations of active and inactive receptors, and as sticks in the insets. Non-polar hydrogens are omitted for sake of clarity.

In the present work, we elucidate the activation mechanism of the adenosine A2A receptor (A2AR), a class A GPCR, providing structural and energetics information on all the functional conformations assumed by the receptor in different ligated conditions. In particular, we performed extensive MD simulations and enhanced sampling calculations that allowed disclosing the free energy landscapes of the (de)activation process of A2AR i) in the *apo* form; ii) in complex with the full agonist 5′-N-ethylcarboxamidoadenosine (NECA); and iii) in complex with the compound ZM241385 (ZMA), whose intrinsic activity (antagonist *vs* inverse agonist) is still debated.^51–57^ In each system, we have identified at atomistic resolution the lowest energy - hence most probable - states assumed by A2AR, rationalising the diverse pharmacological activity of the investigated ligands by elucidating how they affect the receptor activation free energy landscape. In particular, the binding of ZMA locks the receptor in the inactive state, while the agonist NECA predisposes A2AR for the active conformation competent for G protein binding, with the definitive activation of the receptor - and stabilisation of the active form - occurring only after the G protein coupling. Importantly, we disclose two structures of A2AR in the *apo* form that were not previously resolved and connect to the NMR and fluorescence data reported for this receptor.^11,55–60^ One is similar to the inactive state experimentally found in presence of inverse agonist ligands. The other one corresponds to a novel receptor conformation that we named *pseudo-active state* (pAs). This structure is characterized by a distinctive arrangement of the connector region and in particular TM6, with a state-specific orientation of the “activation microswitches” amino acids, such as W^6.48^ (Ballesteros-Weinstein numbering used)^61^ (“toggle switch”), the E^6.30^/DRY^3.49-51^, the P^5.40^I^3.40^F^6.44^ and the NPxxY^7.49-53^ motifs (Fig. 1).^46,62–65^ Among these, the class A conserved residue E^6.30^ plays a leading role during A2AR dynamics, determining the TM6 rotation necessary for receptor activation. In particular, E^6.30^ works as the key “activation” switch by interacting with R^3.50^ in the A2AR inactive state - forming the characteristic “inactivating ionic lock” (IIL) found in many GPCRs^62,63^ - while in the active state it engages a salt bridge with R^5.66^. The phylogenetic conservation of R^5.66^ and E^6.30^ in class A GPCRs prompted us referring to the E^6.30^/R^5.66^ interaction as the receptor “activating ionic lock (AIL)”. Interestingly, in the newly identified pAs, A2AR is able to couple β-arrestin 1 over G proteins, enlightening possible routes for receptor desensitization and G protein-alternative cellular pathways.

Our simulation protocol is generalisable and can be applied to study the activation of any GPCR, resulting a precious tool for biased signaling studies. An explanatory movie of the A2AR activation mechanism is available as Supplementary Movie 1 and at https://www.youtube.com/watch?v=5FrtoZutSa0.

The PDB structure of the A2AR pAs is reported in the Supplementary Materials and at https://www.pdbdb.com/, providing an unprecedented structural basis for the design of A2AR ligands with therapeutic potential for cancer, inflammatory, cardiovascular, Parkinson’s and Alzheimer’s diseases.^66–71^.

## Results

### Effect of ligand binding and G protein recruitment on A2AR conformational dynamics

The A2A receptor (A2AR) can assume a large number of conformations ranging from the active state,^72,73^ bound to agonist and G protein, to the inactive state, typically stabilised by antagonist or inverse agonist binding.^64,74^ In order to investigate how the diverse ligands and G protein influence the receptor dynamics, we first performed a series of extensive all-atom molecular dynamics simulations on differently ligated forms of A2AR. In particular, starting from the experimental structures of the active and inactive A2AR,^72,75^ we prepared seven distinct simulation systems in which the GPCR is coupled with pharmacologically diverse ligands - the agonist NECA, the antagonist/inverse agonist ZMA - and the miniGs heterotrimer (Gs) in all possible combinations (Table 1). The A2AR was embedded in a mixed POPC-cholesterol (7:3 ratio) membrane environment, and each system was simulated in explicit solvent for 5 μs, resulting in a total simulation time of approximately 35 μs.

**Table 1.**
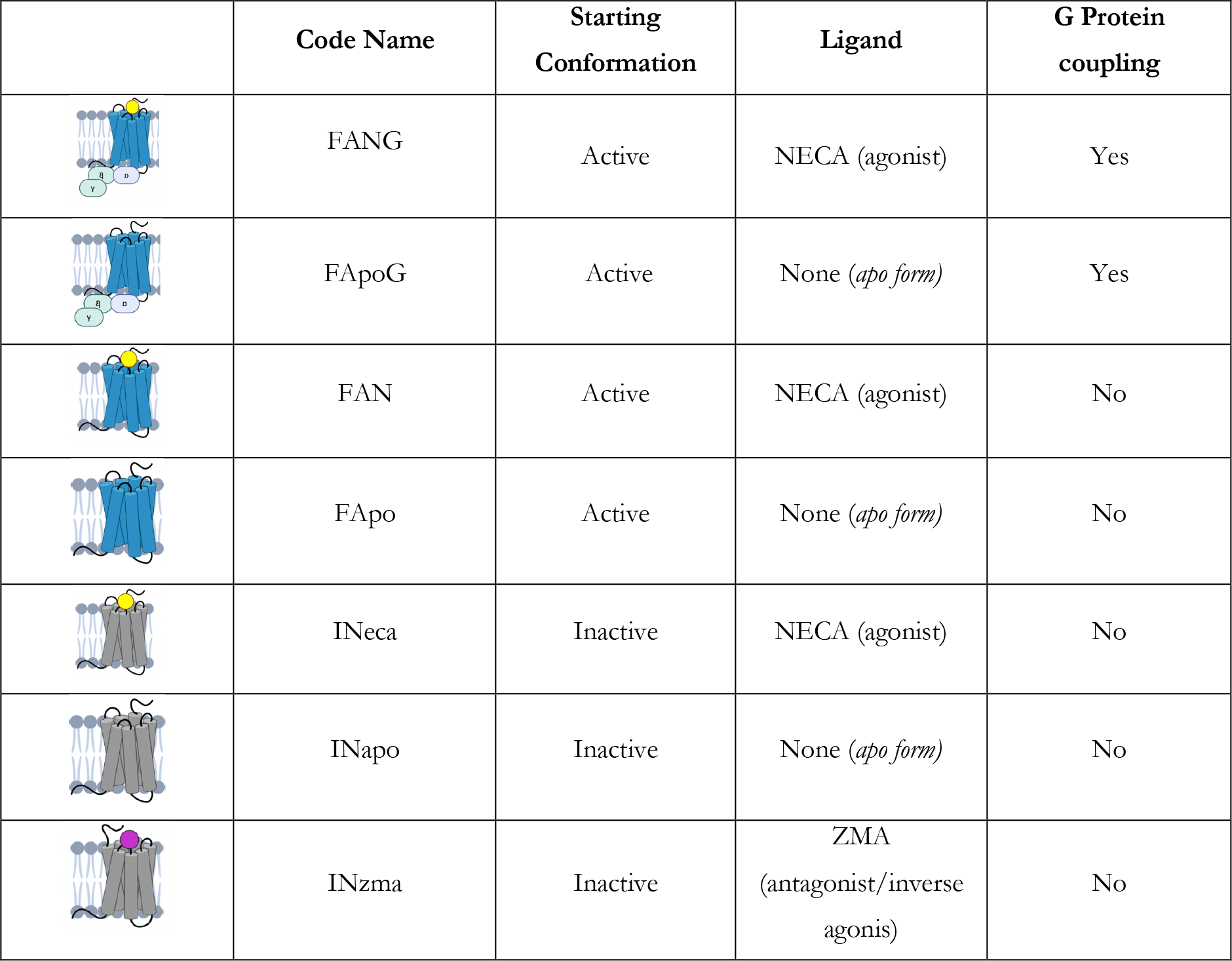
Summary of the simulated systems

|  | Code Name | Starting Conformation | Ligand | G Protein coupling |
| --- | --- | --- | --- | --- |
|  | FANG | Active | NECA (agonist) | Yes |
|  | FApoG | Active | None (*apo form*) | Yes |
|  | FAN | Active | NECA (agonist) | No |
|  | FApo | Active | None (*apo form*) | No |
|  | INeca | Inactive | NECA (agonist) | No |
|  | INapo | Inactive | None (*apo form*) | No |
|  | INzma | Inactive | ZMA<br>(antagonist/inverse agonis) | No |

Our simulations allowed elucidating the stabilizing effect of orthosteric ligands and G protein on their binding sites, the OBS and the IBS, respectively. In the OBS, higher RMSD fluctuations are generally observed for the *apo* systems (1.4 ± 0.2 Å, 1.2 ± 0.3 Å and 1.4 ± 0.2 Å for *FApo, FApoG*, and *INapo*, respectively) compared to the *holo* ones (0.9 ± 0.1 Å, 0.9 ± 0.1 Å and 1.1 ± 0.2 Å for *FAN, FANG*, and *INzma*, respectively) (Supplementary Fig. S1). This phenomenon can be ascribed to the drug-protein interactions engaged by NECA and ZMA, which stabilize the OBS side chains conformations and consequently reduce the backbone fluctuation. Only the *holo Gs-uncoupled* system *INeca* shows relatively high RMSD values (1.36 ± 0.23Å) of the OBS. This is likely due to induced fit effects of the agonist NECA on the inactive form of A2AR, which is not the natural host conformation of the receptor for this ligand. In fact, as no experimental structure of inactive A2AR in complex with the agonist NECA is available, the starting state of the *INeca* simulation was obtained from manually docking the agonist into the *inactive* experimental structure of A2AR (see Methods for details). The purpose of this simulation is to investigate how the ligand influences the receptor conformational dynamics starting from the inactive state and if it could drive the receptor towards the active state in absence of G protein.

Similarly to the ligands at OBS, Gs stabilises the IBS. In fact, the miniGs heterotrimer engages strong and specific interaction with IBS residues, locking A2AR in its active state, as shown by the RMSD comparison (Supplementary Fig. S2) between the two *coupled* systems (1.7 ± 0.4 Å and 1.7 ± 0.4 Å for *FANG* and *FApoG*, respectively) with the corresponding *uncoupled* ones (1.9 ± 0.5 Å and 3.1 ± 0.8 Å for *FAN* and *FApo*, respectively).

Interestingly, allosteric communications between the OBS and the IBS have been found looking at the motion of the intracellular receptor region in the *holo Gs-uncoupled* systems (1.9 ± 0.5 Å and 1.8 ± 0.2 Å for *FAN* and *INzma*, respectively). In fact, in such cases the RMSD of the IBS is lower, in terms of both values and fluctuations, compared to the two *apo Gs-uncoupled* trajectories (3.1 ± 0.8 Å and 2.2 ± 0.4 Å for *FApo* and *INapo*, respectively). This result suggests that the A2AR’s intracellular portion can be stabilised by the binding of a specific ligand to OBS, in addition to the stability given by the binding of transducers at IBS. In this perspective, we investigated the presence of correlated A2AR inter-helical motions in the different simulated systems by computing a Pearson Coefficient (PC) matrix (Figure 2A, Supplementary Fig. S3). Interestingly, this analysis confirmed that the overall receptor dynamics is strongly reduced when either the orthosteric ligand or G protein - or both - are bound to the GPCR. Indeed, looking at the PC maps (Figure 2A, Supplementary Fig. S3) few spots at positive (blue spots, PC > 0.5) or negative (red spots, PC > 0.5) correlation values are found in all the *holo/coupled* systems, whereas significant inter-helical communication areas were observed in the *apo-uncoupled* systems *INapo* and *FApo*. Particularly, among all the simulated systems, *FApo* shows the largest conformational changes with a tight coupling (Figure 2A, Supplementary Fig. S3) between the fluctuations of TM6 and the intracellular parts of TM1-TM2 and TM3-TM4, which, in turn, are mutually anti-correlated. These data indicate a rearrangement of the receptor, especially at intracellular level, in line with the RMSD values computed for the IBS (Figure S2), the receptor’s TMs and the connector regions (Supplementary Fig. S4 and S5, respectively).

**Figure 2.**
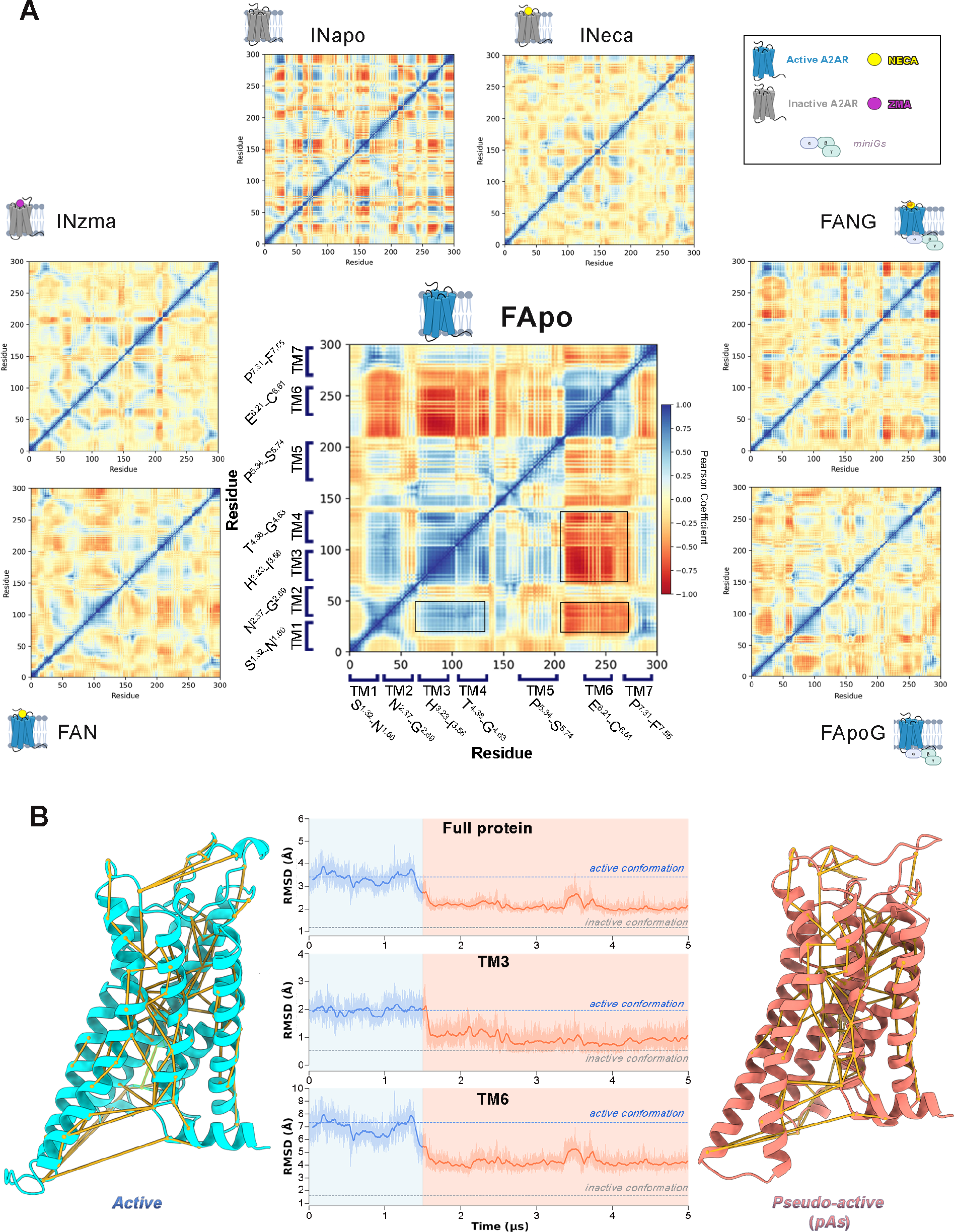
Active to pseudo-active transition. A) Pearson Coefficient Matrices computed for the protein Ca atoms over the seven simulated systems. B) RMSD plots computed with respect to the inactive A2AR conformation for the entire protein, and for both the TM3 and TM6 helices (Ca atoms) over the FApo trajectory. The tridimensional structures of the starting full active conformation and of the newly identified pseudo-active state are shown as cyan and coral cartoon, respectively. The most relevant Protein Structure Network (PSN) metapath of the observed transition are highlighted as yellow links.

Indeed, within ∼1.6 μs the *apo G protein uncoupled* A2AR undergoes a major conformational rearrangement of TM3 and TM6, leaving its starting active state and reaching a conformation intermediate between the active and inactive states (Fig. 2B). Such large-scale motion is characterised by two main events (See Supplementary Movie). In the first one, the two helices move in opposite directions along the axis perpendicular to the membrane plane (*z*), with TM3 shifting downward (intracellularly) and TM6 upward (extracellularly) of ∼3 Å. In the second one, the intracellular segment of TM6 approaches the centre of the TM bundle, assuming a state-specific tilted conformation (Fig. 2). This represents the final state of the receptor that is stable for the rest of the simulation time, longer than 3.4 μs (Supplementary Fig. S6). The newly identified A2AR state, hereafter referred to as *pseudo-active state* (pAs), has never been characterised before and its remarkable structural and energetic stability prompted us to deeply analyse this receptor conformation in the following section. The PDB structure of the A2AR pAs is reported in the Supplementary Materials and at https://www.pdbdb.com/, whereas an explanatory movie of the A2AR activation mechanism is available as Supplementary Movie 1 and at https://www.youtube.com/watch?v=5FrtoZutSa0.

### Structure of A2AR pAs

The tridimensional structure of the newly identified A2AR pAs, corresponding to the most sampled receptor conformation in the pAs state, is rather different from the reported inactive (ZMA-bound, PDB code: 4UM9) and active (NECA and G protein-bound, PDB code: 5G53) states of A2AR, with RMSD values computed for the backbone Ca atoms of 2.52 Å *vs*. 3.28 Å, respectively. A closer inspection of the structure reveals the key residues that stabilise this receptor conformation as transition intermediate between the active and inactive form. Such residues were identified by analysing the A2AR conformations collected during the *FApo* MD simulation through a Protein Structure Network (PSN) model that is able to assess time-related residue-residue interactions (Figure 2B, Supplementary Fig. S3 and Methods for details). Special attention was dedicated to the analysis of residues known as “activation microswitches” in Class A GPCRs^46,62–65^. For the sake of clarity, the following discussion is organised treating separately the three main structural components of GPCRs: the OBS, the connector and the IBS regions. *Orthosteric Binding Site (OBS)*. In this region, pAs is more similar to the inactive conformation than the active one. Proof of that is the lower RMSD values computed for the OBS residues in pAs with respect to those of the inactive and the active state, 0.9Å and 1.25 Å, respectively. This evidence is further confirmed by analysing the conformations of the OBS residues known to be involved in receptor activation such as V84^3.32^, T88^3.36^, S277^7.42^ and W246^6.48^.^72,73,76–78^ In fact, such amino acids occupy a very similar position to that observed in the inactive receptor (Fig. 3A’), while major differences occur with respect to the active conformation (Fig. 3A). In more detail, the sidechains of T88^3.36^ and S277^7.42^ in pAs are oriented outwards the binding site if compared to their position in the experimental active A2AR structures (Fig. 3A). In fact, in the latter structures these two residues engage polar interactions with the agonists’ ribose ring, which are instead missing when the receptor is either in its *apo* form or bound to antagonists/inverse agonists.^72,73,77–79^Also the rotameric state of V84^3.32^ and W246^6.48^ (“toggle switch”) is more similar to the inactive conformation than the active one in which the agonist’s ribose ring shifts V84^3.32^ and W246^6.48^ towards an outward and downward conformation, respectively (Fig. 3A-B).

**Figure 3.**
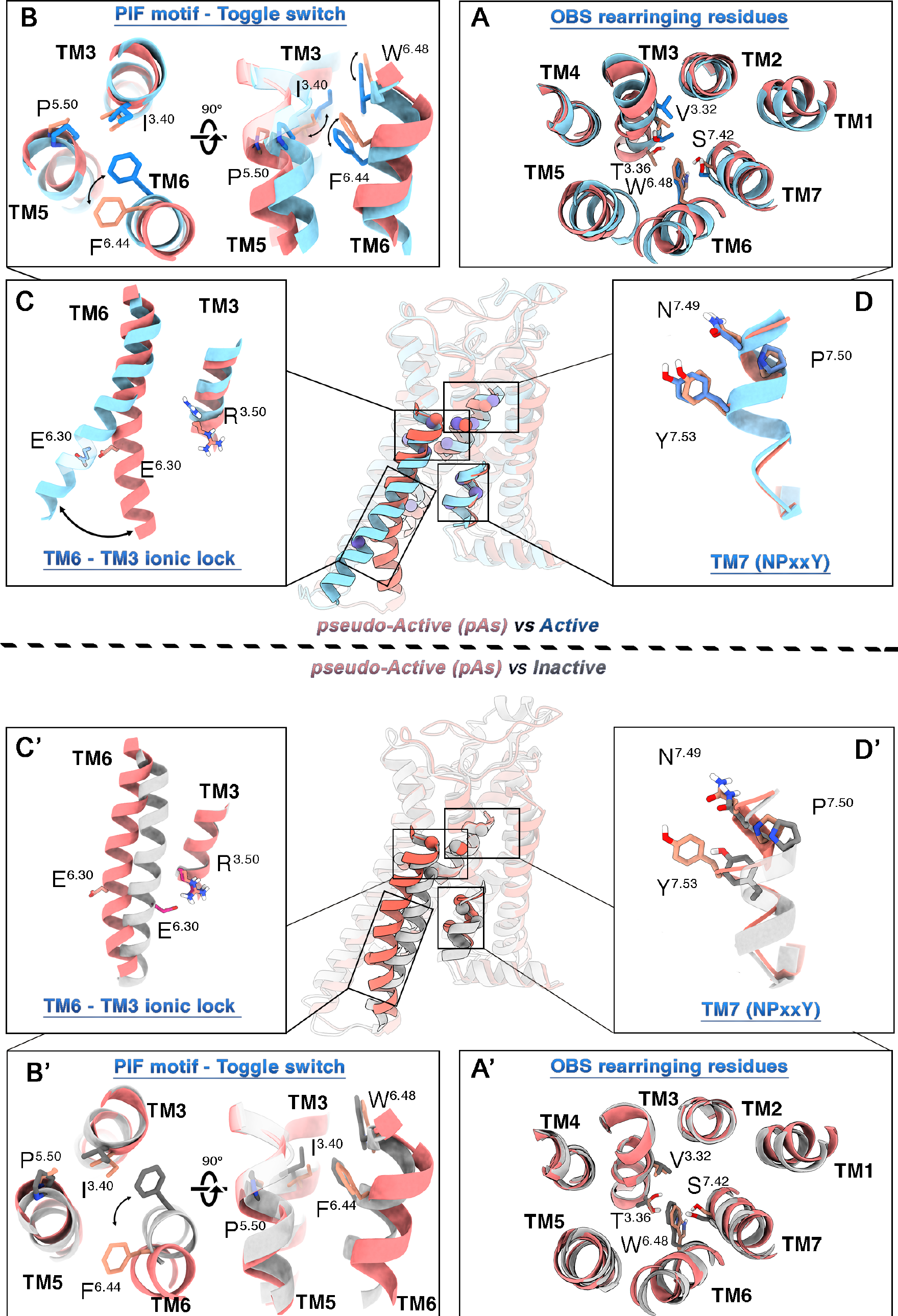
Atomistic details of the newly discovered A2AR pAs. The conformations of the main activation microswitches belonging to the A2AR pAs (salmon) were compared with both the active (PDB code: 5G53, color: light blue, upper panel) and the inactive (PDB code: 3PWH, color: silver, lower panel) A2AR states.

*Connector Region*. At variance with the OBS, the conformation assumed by the pAs connector region is state specific, dissimilar from both the active and the inactive states - RMSD values of 2.3Å and 2.4 Å, respectively - and is characterized by a distinctive orientation of the PIF motif. Comparing pAs to the active A2AR, TM6 shows an upward movement along the *z* axis and a counterclockwise rotation, which orients the F242^6.44^ sidechain outwards with respect to the active and inactive structures (Fig. 3B-B’). On the other hand, TM3 is shifted downwards, with I92^3.40^ assuming its inactive state position (Fig. 3B’). This is an unprecedent finding in the context of the A2A (de)activation mechanism. In fact, in all the structural studies reported so far, TM6 was found to move inward and rotate clockwise when passing from the active to the inactive A2AR.^64,72,73,77–79^ Instead in pAs, TM6 moves in the opposite direction and is rotated counterclockwise to facilitate the vertical motion of TM3 and the onset of the deactivation process.

*Intracellular Binding Site (IBS)*. The intracellular portion of pAs is characterised by a bending of TM6. Specifically, the cytoplasmic end of TM6 is bent towards the centre of the TM bundle (Fig. 2,3C’), reducing the volume of the IBS to values comparable to that of the inactive A2AR X-ray structure (Supplementary Fig. S7). However, two major differences arise by comparing the pAs and the inactive-state IBSs. First, the middle-lower portion of TM6 (residues 229-237) is slightly shifted outwards (Fig. 3C’). This might be due to the steric hindrance of the NPxxY motif at TM7 in the pAs. In fact, such motif assumes a position very similar to that of the active A2AR (Fig. 3D), directly interacting with TM5 through the known water-bridged H-bond between Y288^7.53^ and Y197^5.58^.^65^ The second relevant feature is that TM6 is rotated about 40°-50° counterclockwise with respect to the inactive state (Fig. 3C’). This conformation is stabilised by a salt-bridge interaction between R205^5.66^ of TM5 and E228^6.30^ of TM6 (Fig. 4A). Noteworthily, a similar interaction was found in the active state structures of the Rhodopsin receptor – where R^5.66^ is mutated into K^5.66^ (PDB: 3CAP and 3DQB)^34,80,81^ – and in the A3 adenosine receptor by in silico studies.^82^ The results of our simulations show that such ionic interaction is very stable in all the A2AR structures with TM6 in an active-like rotameric state (frequency of occurrence > 95%, Fig. 4B), whereas it is lost in all the A2AR inactive conformations (Fig. 4B). In the latter, E228^6.30^ is oriented toward the inner part of the IBS and interacts with R102^3.50^ of the DRY motif, forming the “inactivating ionic lock” (IIL). Interestingly, R205^5.66^ and E228^6.30^ are highly conserved among the Class A GPCRs (37% and 35%, respectively, Fig. 4C), prompting us to propose the E228^6.30^-R205^5.66^ salt-bridge as the “activating ionic lock” (AIL), *alter ego* of the IIL, whose loss facilitates the clockwise rotation of TM6 and in turn the deactivation process.

**Figure 4.**
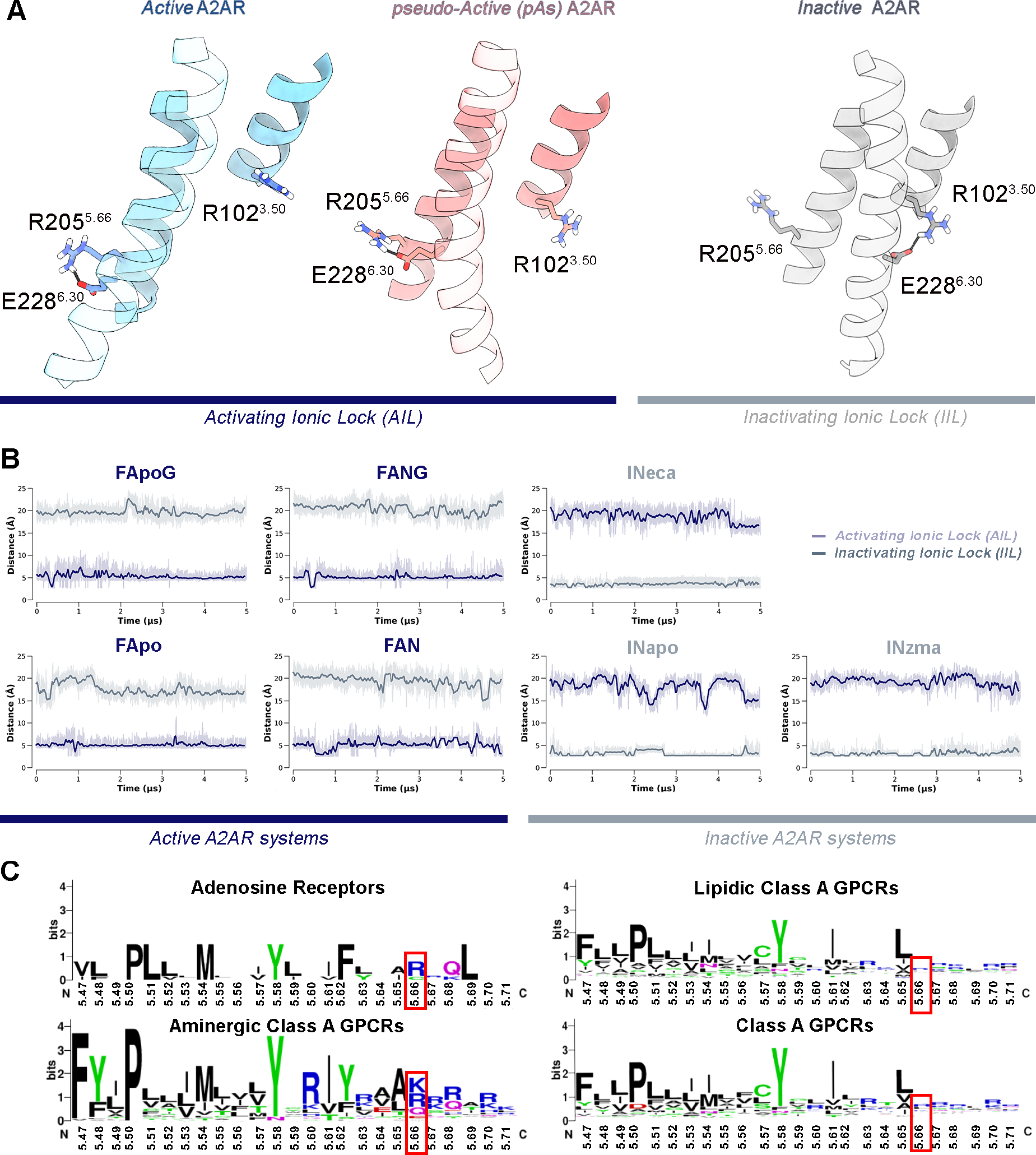
A) 3D representation of the activating ionic lock (AIL) and inactivating ionic lock (IIL) in the A2AR’s active, pseudo-active and inactive states. The helices TM3, TM4 and TM6 are depicted as cartoon while residues E288^6.30^, R201^5.66^ and R98^3.50^ are highlighted as sticks. B) Plots of the AIL (distance between the cγ atom of E288^6.30^ and the cζ atom of R201^5.66^) and IIL (distance between the cy atom of E288^6.30^ and the cζ atom of R102^3.50^) along the seven MD systems. C) LOGOs analysis of residues 181^5.47^-206^5.71^ of TM5. The conservation percentages for basic aminoacids at position 5.66 in the four investigated subsets (Adenosine, Aminergic, Lipidic and entire Class A) of Class A GPCRs are respectively: 75%, 68%, 38% and 37%.

The role of the new pAs along the A2AR transition from the active to the inactive state was further investigated by analysing the MD simulations of the *apo GP-uncoupled* A2AR. Our results show that within the MD µs timescale, pAs can be reached from the active state (*FApo* system), but not from the inactive one (*INapo* system), which is instead stable throughout the simulation (Supplementary Fig. S4). Furthermore, pAs is a very stable, long-lasting state with a residence time longer than 3 μs (Supplementary Fig. S6). This finding indicates that pAs is a metastable, intermediate conformation between the active and inactive ones, which are separated by a relatively large energy barrier that is unlikely to be crossed within µs (simulation) timescale. Our results further show that in absence of the orthosteric ligand and G protein, the receptor can also assume a stable inactive state, in agreement with NMR and fluorescence-based studies^11,55,83^ that found the *apo* A2AR in a state similar to the antagonist/inverse agonist-bound conformation. In order to observe the transition between the active and inactive states, it is necessary to accelerate the sampling and overcome the timescale limitation of standard MD simulations. This is possible by employing enhanced sampling calculations based on Well-Tempered Metadynamics (WT-MetaD)^84,85^ combined with Path Collective Variables (PCVs) (See Methods for details).^86^ Particularly, PCV is a dimensionality reduction approach suitable to describe largescale protein motion, taking into account multiple degrees of freedom of a system for which the terminal states are known, as in the A2AR case. The process under investigation is thus accelerated by applying a bias potential along the PCV that is defined as a path composed by a sequence of intermediate frames connecting the two terminal system’s conformations (A2AR active and inactive states in our case). We note that during the PCV WT-MetaD calculations the receptor can explore conformations even different from the original path, thus making the results independent from the choice of the original path. PCV-MetaD calculations have been successfully employed by us and other groups to study large scale and long timescale conformational changes in different protein systems.^87–93^ In the case of A2AR, the path is defined considering pAs as an intermediate state between the terminal active and inactive state, and including the residues involved in the conformational transitions observed during the multi-microseconds MD calculations (see Methods for details).

### Activation Free Energy Landscape of A2AR

The entire (end-to-end) activation and deactivation process of A2AR was investigated by means of PCV-MetaD in three different systems: i) *apo A2AR; ii) NECA-bound* A2AR; and iii) *ZMA-bound* A2AR. In all of them, the sampling was enhanced by adding a bias potential on two PCVs, containing all the inter-residue contacts involved in the conformational receptor transitions observed in the previously discussed MD calculations (see Fig. S8 and Methods for the details). In particular, the first PCV (_*ACT*_P) is defined as an RMSD matrix describing the geometric distance of the backbone atoms involved in the active-to-inactive receptor transition (see Supplementary Table S1 and Fig. S8A). The second PCV (_*TM6*_P) is instead defined as a contact map (CMAP) between residues characterising the rotation of TM6, clockwise from active to inactive (see Supplementary Table S2 and Fig. S8B). The three free-energy calculations reached convergence at different simulation times: 2.4 μs for *apo-*A2AR, 3.5 μs for *NECA-bound-*A2AR, and 3.6 μs for *ZMA-bound*-A2AR, for a total of 9.5 μs of enhanced sampling simulations (Fig. S9-11). Considering the acceleration factor computed during the MetaD calculations - 10^9^-10^10 -^ (See Methods for detail), we could reasonably estimate that the observed receptor activation and deactivation process occur on a minute timescale. For each system, at the end of the simulation we computed the activation free-energy surface (FES) as a function of two CVs, _*ACT*_P.s and _*ACT*_P.z (See Methods for details). The first CV (_*ACT*_P.s) describes the receptor exploration of the different states forming the transition path from the active to the inactive state, whereas the second CV (_*ACT*_P.z) defines the distance as MSD of the sampled conformations from the reference path (Fig. 5A). As previously introduced, using PCV-MetaD the systems can explore conformations even distant from the reference path that, in such a case, would have high _*ACT*_P.z values. Interestingly, in the three systems all the low energy minima - hence most probable receptor states - have low _*ACT*_P.z values (< 0.05 nm^2^). This result indicates that the reference path employed in WT-MetaD calculations well represents the low energy transition path from active to inactive A2AR, leading to a reliable description of the receptor (de)activation process.

**Figure 5.**
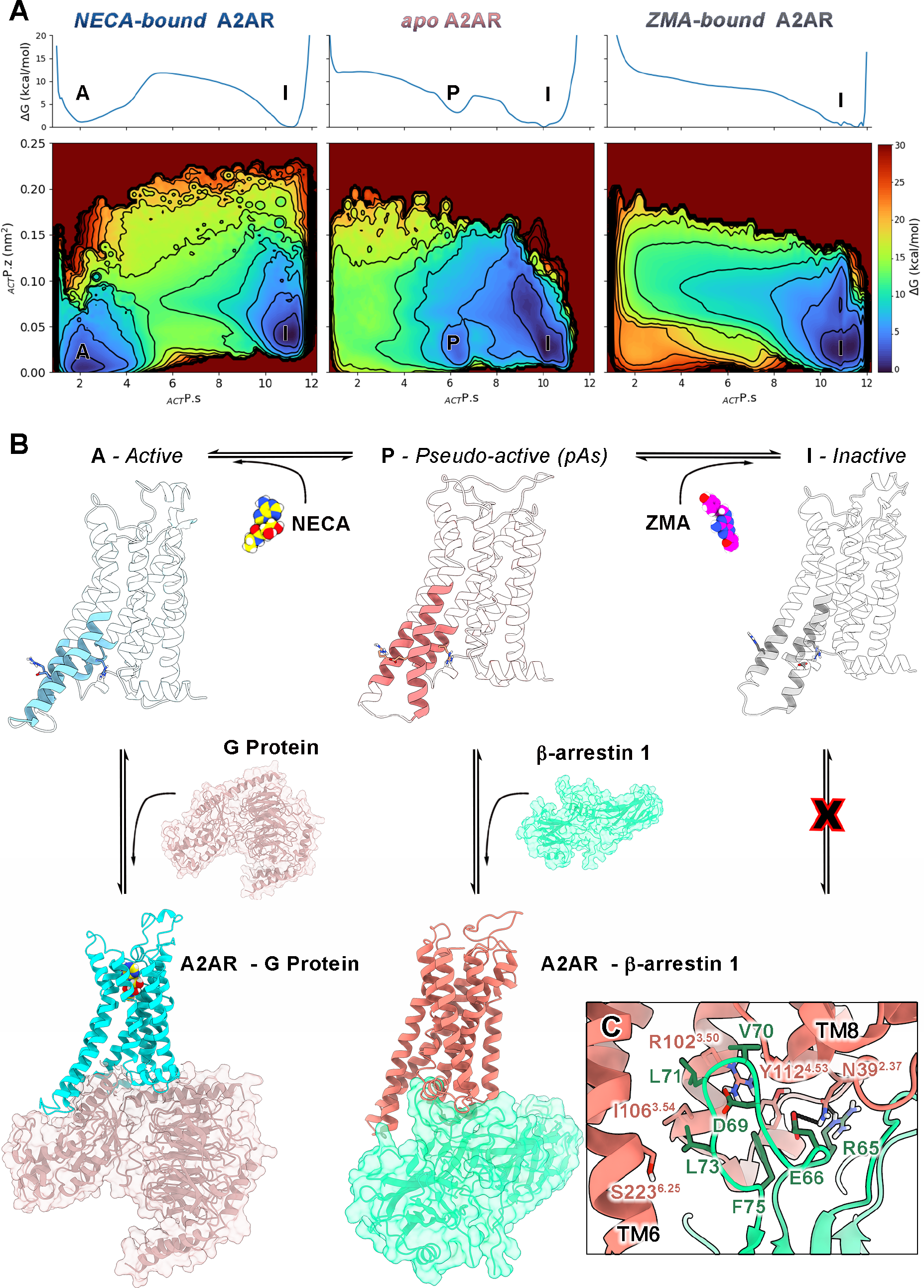
A) Activation Free Energy landscapes of the NECA-bound, apo and ZMA-bound forms of A2AR as a function of the _*ACT*_P.s and _*ACT*_P.z collective variables. Isosurfaces are displayed every 3 kcal/mol. B) Atomistic representation of the A2AR structures corresponding to the main free energy minima (A,P,I) and equilibria interconnecting the same conformations along the activation process. C) Docking-predicted binding mode of βarrestin 1 (green cartoon and sticks) at the pAs IBS (salmon cartoon and sticks).

Comparing the three FESs in Fig. 5, it is possible to assess the effect of ligand binding on the low energy states assumed by the receptor during its active-to-inactive transition. In detail, the *NECA-bound* A2AR is in equilibrium between two energetically comparable states (free energy difference between minima is 0.8 ± 0.5 kcal/mol). The structure representing the energy minimum A corresponds to the active crystallographic state (RMSD value is 1.4 Å computed for the TM helices with respect to PDB code 5G53), whereas the structure representing the energy minimum I is very similar to the inactive receptor state (RMSD value is 1.6Å computed for the TM helices with respect to PDB code 5IU4). This finding indicates that upon agonist binding i) the receptor can reach the active conformation competent for G protein binding; and ii) the inactive form remains the lowest energy receptor state in absence of G protein. Similarly to the *NECA-bound* system, the *apo* A2AR has two lowest energy conformations. While the lowest energy minimum is still represented by the A2AR inactive conformation (state I) (RMSD = 1.3 Å), the second energy minimum, 2.9 ± 0.3 kcal/mol higher than I, is not the active state, but state P that corresponds to the pAs structure previously identified by our unbiased MD calculations (RMSD < 1.2 Å). Therefore, the free-energy calculations confirm the presence of the *pseudo-active* state pAs as metastable intermediate between the active and inactive form of *apo* A2AR. Differently, the presence of ZMA at the OBS shifts the conformational equilibrium toward the inactive state. In fact, the *ZMA-bound* A2AR has one single low energy minimum (state I), corresponding to the crystallographic pose of the receptor in complex with ZMA. Comparing the FESs in Fig. 5, one can see that in *ZMA-bound* A2AR both the active and pseudo-active state are lost in favor of the receptor inactive state. The energetic and structural stability of all the identified minima was further evaluated through µs-long unbiased MD simulations (A, P, and I) (Supplementary Fig. S12).

In order to investigate in more detail the molecular features characterizing A2AR activation, we computed the FES as a function of _*ACT*_P.*s* and _*TM6*_P.*s* (Supplementary Fig. S13). As introduced before, the _*ACT*_P.*s* CV defines the active-to-inactive receptor transition - specifically the outward/inward motion of helices TM5-6-7 -while the second CV (_*TM6*_P.*s*) describes the *around-the-axis* rotation of TM6 (clockwise or counter-clockwise). Looking at these FESs, the highest energy barriers between basins A and I, and between P and I, are found along the _*TM6*_P.s CV (13.4 ± 0.5 kcal/mol and 14.5 ± 0.2 kcal/mol, respectively). Therefore, the TM6 rotation represents a slower degree of freedom than the TM5-6-7 outward/inward motion and can be considered the rate determining step of the (de)activation process. Three hubs of microswitch residues rule the receptor motion and are: i) the salt bridges engaged by E228^6.30^ with R205^5.66^ in the active form (AIL - *activating ionic lock*) and with R102^3.50^ in the inactive one (IIL – *inactivating ionic lock*); ii) the Van der Waals interactions established by L235^6.37^, I238^6.39^, V239^6.40^ with L194^5.55^, L198^5.59^ and F201^5.62^ in the active and pseudo-active states, and with L95^3.43^, I98^3.46^ and I200^5.61^ in the inactive states; and iii) the water-bridged interaction between Y197^5.58^ and Y288^7.53^ in both the active and pseudo-active states. The loss of the latter allows the outward motion of the NPxxY motif characterizing the receptor deactivation.

### The pAs/β-arrestin 1 complex

The discovery of the pAs prompted us to investigate if such receptor state might play a functional role. To this end, we performed protein-protein docking calculations on pAs with the two effector proteins Gs protein (i.e., miniGs) and β-arrestin 1. Our results revealed that the coupling of pAs with G protein is prevented by the steric hindrance of the TM6 intracellular tail (Figure S14A), which significantly reduces the A2AR IBS’s volume in pAs (see Supplementary Fig. S7). Interestingly, a high binding score is instead computed between pAs and β-arrestin 1, consistent with previous structural investigations^94–96^ showing that a reduced volume at the intracellular pocket favors the interaction with β-arrestins. The resulting binary complex exhibits very similar protein-protein interactions to the Cryo-EM structure of the β1-adrenoreceptor (β1AR) coupled with β-arrestin 1. Examples are the polar contacts between A2AR’s R102^3.50^ and Asn39^2.37^ (R139^3.50^ and Asn74^2.37^ in β^1^AR) with β-arrestin’s Glu66 and the hydrophobic interactions formed by A2AR’s Ile106^3.54^ (Ile143^3.54^ in β^1^AR) and β-arrestin’s Leu71 and Leu 73 (Figure 5C and S14). These findings indicate that pAs is an energetically stable state of A2AR that might play a role in β-arrestin-mediated activation of specific cellular pathways.

## Discussion

In this study, we have provided a thorough structural and energetics characterisation of the activation mechanism of the adenosine G protein coupled receptor A2AR. Specifically, the *apo*, the *agonist-bound* and the *inverse agonist-bound* forms of the receptor have been investigated. Among these the A2AR *apo* form is particularly interesting as this represents the basal functional state of the receptor for which no experimental structure has been reported so far. In this condition, the receptor can assume the inactive form, as also outlined by previous NMR and fluorescence studies^11,55,83^, in equilibrium with a pseudo-active state pAs, which is here disclosed for the first time (*P-I equilibrium*, Figure 5B). In the pAs structure the receptor presents a mix of molecular features of the active and inactive state. The most important one is the salt bridge interaction established between E228^6.30^ and R205^5.66^ - largely conserved within class A GPCRs - which stabilises the TM6 active-like orientation, forming what we have defined activating ionic lock (AIL). The newly identified state pAs is able to couple β-arrestin 1, showing a binding mode very similar to the experimental one found between β^1^AR and β-arrestin 1 (Figure 5C and S14). This finding opens novel opportunities for modulation of the A2AR activity, especially in terms of receptor desensitization and activation of G protein-alternative pathways mediated by β-arrestins.^97–99^ For instance, pAs might be targeted to identify novel A2AR biased ligands using standard and advanced drug discovery campaigns that employ AI-based algorithms like graph neural networks and geometrical deep learning to increase the hit-discovery success rate. In addition, pAs might be helpful in elucidating the yet unclear molecular aspects of β-arrestins coupling to A2A, including phosphorylation at the A2A C-terminus.^97,100,101^

When A2AR is bound to the agonist NECA without G protein, the receptor is found in two most probable states (*A-I*, Figure 5B). The first one corresponds to the inactive structure of the receptor, in agreement with the reported X-Ray and Cryo-EM agonist-bound structures of A2AR^77–79,102^ and as also found for the β^2^-adrenoreceptor by structural and computational studies^103,104^. At this regard, it is worth noting that the experimental agonist-A2AR complexes (*GP-uncoupled*) are loosely defined as intermediate states, however all of them have TM6 rotated clockwise in the inactive form and the intracellular region of the receptor close to the inactive state. In fact, plotting the position of the experimental agonist-bound A2A structures^77–79,102^ onto the FES of the activation mechanism, they all result very close to the inactive state I (Figure S15). The second low-energy state instead corresponds to the receptor active form, characterised by the counterclockwise rotation of TM6 and the presence of the AIL. This indicates that the agonist binding - even in absence of G protein - induces conformational changes in A2AR leading to the loss of pAs in favour of the active form (*A*), which is competent for the G protein binding. Our finding rationalises the ^19^F-NMR data of *agonist-bound* and *apo* A2AR reported by Huang et al.^55^ (Figure S16) which show in both cases the A2AR in equilibrium between multiple states, among which one is always represented by the inactive state, whereas the others (i.e. active and pAs) are differently stabilised by the presence of the agonist. Furthermore, our results suggest that the *active-inactive (A-I) equilibrium* established upon agonist binding is functional for the definitive activation of the receptor - and stabilisation of the active form - occurring only after the G protein recruitment at the receptor IBS.

Finally, when the receptor is bound to ZMA, the basal *P-I equilibrium* is shifted towards the inactive conformation, which is the only low energy state present (Figure 5A). We note that the intrinsic pharmacological activity of ZMA is debated, with studies reporting this ligand as antagonist,^51–53^ while more recent works attribute to ZMA an inverse agonist activity.^55–57^ Based on the widespread knowledge that an antagonist competes with the agonist for binding without altering the receptor basal activity, while an inverse agonist also reduces the basal activity, our results indicate that ZMA works as inverse agonist since it locks the A2AR in the inactive form. In fact, if ZMA worked as antagonist, it should have not altered the two-states equilibrium of the A2AR *apo* form.

Our study provides unprecedented structural and energetics insight into A2AR activation mechanism, revealing a novel receptor pseudo-active state that could be further investigated in biased signaling studies and targeted by drug design campaigns to develop A2AR biased ligands. Our simulation protocol is generalisable and can be applied to study the activation mechanism of any GPCRs and to predict the intrinsic activity of ligands based on their effect on the receptor conformational dynamics, resulting a precious tool for investigations on GPCRs activation and drug design.

## Methods

### Systems setup and Unbiased Molecular Dynamics

According to the best resolution criterion, the starting conformations of the active and inactive A2AR have been taken respectively from the 5G53^72^ and 5IU4^75^ X-ray structures, whereas the coordinates of the mini G_s_ heterotrimer were extracted from the cryo-EM structure 6GDG^73^ (after the alignment of the GPCR section with 5G53). The *active GP-uncoupled* (*FAN, FApo*) and the *apo* (*FApo, FApoG* and *INapo*) systems were obtained by removing the engineered G protein or the co-crystalized orthosteric ligands (or both) from the original pdbs (5G53^72^ and 5IU4^75^). To prepare the *INeca* system, we first aligned the agonist- and antagonist-bound A2AR, and then replaced the antagonist ZMA in the inactive receptor conformation (pdb: 5IU4^75^) with the agonist NECA. Any mutation present in the starting pdbs was converted to its wild-type form. Before MD simulations, the receptor first N-terminal (S5) and the last C-terminal (S305) residues were capped with acetyl and N-methyl groups, respectively. Where residues of the intra-cellular loops were missing, they were added and refined using Prime.^105,106^ The receptor conformations were prepared using the Protein Preparation Wizard tool, implemented in the Maestro Suite 2021.^107^ Correct bond orders were assigned, missing hydrogen atoms added and all the water molecules deleted from the receptor structure. Then, a prediction of the side chains ionization and tautomeric states was performed using Epik.^108,109^ Finally, the receptor hydrogen-bonding network was optimized, and the position of all the hydrogens minimized. Each optimized receptor conformation was then embedded in a 105 Å x 105 Å (along x and y axes) pre-equilibrated 1-palmitoyl-2-oleoylphosphatidylcholine (POPC)—cholesterol (7:3 molar ratio) bilayer and solvated using the TIP3P water model with the aid of the membrane-builder tool of CHARMM-GUI.org (http://www.charmm-gui.org). The ff14SB and lipid17 Amber force fields were used to parametrize the protein and the lipids, respectively.^110^ As for the GPD molecule and for the two orthosteric ligands, namely NECA and ZMA, different force fields were used based on the distinct chemical nature of the three compounds. Specifically, a combination of Amber OL3^111^ and gaff^112^ (generalized amber force field) force fields parameters were adopted for GPD and NECA, while the gaff^112^ alone was used to treat ZMA. Their atomic partial charges were instead computed using the two-stages restrained electrostatic potential (RESP)^113,114^ fitting procedure implemented in Antechamber.^115^ The electrostatic potentials (ESP) were first calculated through the quantomechanical package Gaussian 16.^116^ Specifically, the adopted protocol included a double step geometry optimization procedure at Hartree–Fock level of theory: i) a preliminary calculation with the 3–21G basis set, followed by ii) a more accurate procedure with the 6-31G* basis set, after which the ESP potentials were computed. The topology files of the systems were obtained with the tleap program of AmbertTools20^117^ and then converted into the Gromacs format by the means of ParmEd. The GROMACS 2020.6^118^ code was used to perform the simulations. A cutoff of 12 Å was used for short-range interactions. The long-range electrostatic interactions were computed through the particle mesh Ewald method^119^ using a 1.0 Å grid spacing in periodic boundary conditions. The non-iterative LINCS^120^ algorithm was applied to constraint bonds, which allowed using a 2 fs integration time step. To solve all the steric clashes, each system underwent 30,000 steps of steepest descent energy minimization in three phases. In the first one, the system heavy atoms were kept fixed to relax only the hydrogens and the water molecules; during the second stage also the lipidic bilayer was released; and in the third step all the atomic positions were minimized. Then, each complex was equilibrated and heated up to 300 K, alternating NPT and NVT cycles (for a total of 30 ns) with the Berendsen coupling bath and barostat,^121^ while applying gradually decreasing harmonic constraints on the heavy atoms of the membrane, protein, and ligands.

For each system, production runs of 5 μs were performed in the NPT ensemble; the pressure of 1 atm and the temperature of 300 K were kept constant with the stochastic velocity rescaling^122^ and the Parrinello-Rahman^123^ algorithms, respectively.

### Cross-correlation analysis

Cross-correlation analysis (or Pearson-correlation coefficient analysis) was used to assess the correlated motions between pairs of residues in the seven simulated MD systems (Table 1). An in-house code was employed to calculate the Pearson coefficients matrices according to the following formula:

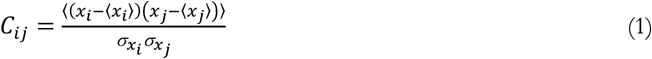

where the numerator is the covariance between two variables, ***x***_***i***_ and ***x***_***j***,_ while ***σ***_***i***_ and ***σ***_***j***_ are the standard deviations of each variable. The normalization obtained dividing the covariance by the product of the standard deviation of the variables allows having values ranging between -1 and +1. The variables represent the Cα atoms’ positional vectors and the Pearson correlation coefficients have been evaluated between any pairs of Cα atoms.

### Protein Structure Network Analysis

Network parameters such as hubs, communities, and structural communication analyses were obtained by using the WebPSN 2.0 web-server.^124–126^ The methodology builds the Protein Structure Graph (PSG) based on the interaction strength of two connected nodes:

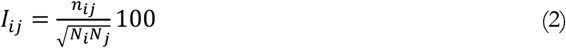

where interaction percentage (***I***_***ij***_) of nodes ***i*** and ***j*** represents the number of pairs of side-chain atoms within a given cut-off value (4.5°Å), while ***N***_***i***_ and ***N***_***j***_ are normalization factors. The interaction strength (represented as a percentage) between residues ***i*** and ***j*** (***I***_***ij***_) is calculated for all node pairs. If ***I***_***ij***_ is more than the minimum interaction strength cutoff (***I***_**min**_) among the residue pairs, then is considered to be interacting and hence represented as a connection in the PSG.

### Well-Tempered Metadynamics with Path Collective Variables

Metadynamics^84^ is an enhanced sampling method in which the simulation is boosted by a gaussian-shaped history-dependent bias potential (V_*G*_), deposited on a selected number of reaction coordinates (i.e. slow degrees of freedom) of the system, usually referred to as collective variables (CVs):

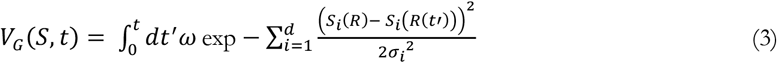

where ***S***_***i***_ is the value of the *i*^th^ CV, ***σ***_***i***_ is the width of the Gaussian function and ***ω*** is the rate at which the bias is deposited. Well-Tempered Metadynamics (WT-MetaD)^85^ is an evolution of the method in which the bias deposition rate ***ω*** is exponentially rescaled over time depending on how much potential has already been added in the same region of the CV phase space, according to following formula:

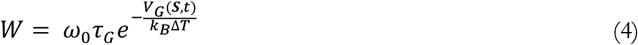

where ***W*** is the Gaussian height, ***kB*** is Boltzmann’s constant, ***ω***_***0***_ is the initial energy rate, ***τ***_***G***_ is the Gaussian deposition stride, **Δ*T*** is the fictitious temperature at which the biased CV (***S***) is sampled and ***V***_***G***_**(*S, t***) is the bias potential accumulated in ***S*** over time ***t***. At the end of a WT-MetaD simulation the deposited bias potential ***V***_***G***_ asymptotically converges to the inverse value of a fraction of the free-energy ***F(S)***:

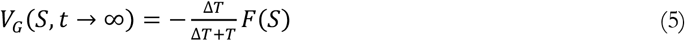

The fictitious temperature ***ΔT*** is the parameter that controls how quickly the Gaussian height is decreased and often is written in terms of the so-called bias factor ***γ = (T + ΔT)/T***. The acceleration factor ***α*** introduced by the underlying metadynamics bias deposited during the simulations was computed as 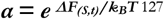 ^127^ using the energetic difference ***ΔF*** calculated between the lowest energy minimum and the highest energy transition state identified in the apo and NECA-bound systems (14.5 kcal/mol and 13.4 kcal/mol, respectively).

The large-scale conformational differences between the crystal structures of the active and inactive A2AR suggest that the transition between these states is highly cooperative and involves a number of degrees of freedom. For this reason, the use of simple geometrical CVs (i.e. a distance or a torsion) might be insufficient both to reproduce the event and to calculate the associated free energy. To overcome this limitation, we employed the path collective variables (PCVs) approach^86^, which has been successfully applied in a number of conformational transition studies.^87–90^ In this dimensionality reduction scheme, two functions are used to characterize the position of a point in a configurational space **R** relative to a preassigned path ***l*** in terms of progress along the path ***s***(**R**) and distance from it ***z***(**R**):

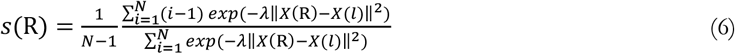

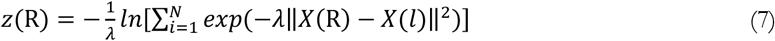

where ***X(R***) is a reduced representation of ***R, X*(*l***) is the same kind of reduced representation of the path ***l, i*** is a discrete index ranging from 1 to ***N***, with ***N*** being the number of conformations selected to build the path. ∥…∥ indicates the metric used to compute the distance between the configurations, which is generally defined in terms of contacts matrix or RMSD. In this work, a preliminary guess of the A2AR activation process was obtained through the unbiased MD system *FApo* (transition from the active to the pseudo-active state) and targeted-MD (transition from the pseudo-active state to the inactive state). The latter was performed applying a constant force (harmonic constant *k =* 4.774 kcal/mol) on a collective variable represented by the RMSD of the protein Ca atoms computed with respect to the experimental inactive conformation of A2AR. The overall trajectory was then used to extract the frames needed for the preassigned paths of two sets of PCVs: P_*ACT*_ and P_*TM6*_.

_***ACT***_***Path Collective Variable*s:** In _*ACT*_PCVs, ***X(R***) is defined as a set of Cartesian coordinates belonging to a subset of atoms. The distance ∥…∥ of each generic configuration ***X(R***) from the path was computed as the root mean square displacement (RMSD) of the subset after optimal receptors alignment by using Kearsley’s algorithm^128^. Notably, the choice of the atoms to be included in the path is far from trivial, in fact, a wrong choice can turn into a loss of performance and additional noise that may affect the calculations. Here, we used the Cα and Cβ of the most important receptor microswitches (PIF, toggle, NPxxY, AIL, IIL) as well as the C and Cy of key residues detected in our unbiased MD (Table S1, Supplementary Figure S8A). The final path consists of 12 frames: 10 frames directly extracted from the preliminary deactivation trajectory (obtained as described above), and two terminal extra-(non-real)-frames. We verified that the obtained set of configurations was equally spaced in the adopted mean square displacement metrics, and the value of λ was chosen so as to be comparable to the inverse of the RMSD between successive frames. The average distance between adjacent frames was 0.74 Å, thus requiring to set λ = 1.22 Å^-2^ in eqs 2 and 3.

_***TM6***_***Path Collective Variables*:** Here, the reduced representation ***X***(***R***) is defined as the CMAP matrix ***C(R***), and the distance ∥…∥ computed as:

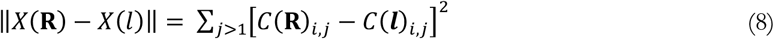

where ***C(R)***_***i***,***j***_ and ***C(l)***_***i***,***j***_ are the elements of the CMAP matrix. A contact between atom ***i*** and ***j*** is defined as

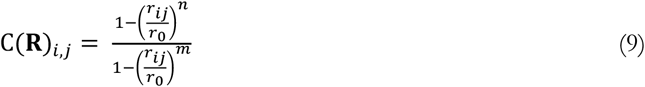

where ***r***_***i***,***j***_ is the distance between the two atoms and ***r***_***0***_ is the typical distance at which the contact is formed (Table S2). The list of the contacts included (Table S2) in the CV was chosen to specifically accelerate the *around-the-axis* rotation of TM6 occurring upon receptor (de)activation. The final path consists of 7 frames. However, The FESs in Fig. S13 and S15 are computed through a reweighting procedure^129^ as a function of path CVs defined by the _*TM6*_PCV used in the production runs with the addition of two terminal extra fames that might represent multiple receptor conformations sampled at the end points. The average distance between adjacent frames was 0.63, thus requiring to set λ = 3.64 according to the same criterion used for S_*ACT*_.

The PLUMED 2.7.1^130,131^ library patched with the Gromacs 2020.6^118^ MD engine was used to perform WT-MetaD simulations on the uncoupled A2AR in its *apo, NECA-bound* and *ZMA-bound* forms. Two-dimensional gaussians were added on the ***s***(**R**) components of _*ACT*_PCVs *and* _*TM6*_PCVs every 2 ps. An initial gaussian height of 0.95 kcal/mol was gradually decreased based on a bias factor γ = 15. The gaussian widths were respectively set to 0.1 and 0.03 for the _*ACT*_P.*s* and for _*TM6*_P.*s* dimensions according to the CVs’ fluctuations observed in the standard MD regime. To limit the exploration of unphysical states during the simulations harmonic restraints were placed on the helicity of TM6, based on the ALPHARMSD variable defined in PLUMED. The bias reweighting procedures used along this work were performed according to the algorithm developed by Bonomi et al.^129^

### Protein-Protein Docking

To evaluate the possible affinity of the newly discovered A2AR pseudo-active state (pAs) toward different intracellular transducers such as mini Gs and β arrestin 1, protein-protein docking were performed with the aid of the HADDOCK 2.4 webserver.^132,133^ Specifically, the minimized structure of the pAS extracted from our MD simulations was used as starting conformation for the A2AR, while the 3D coordinates of the mini Gs and β arrestin 1 were taken from the experimental PDBs 6GDG^73^ and 6TKO^95^, respectively. Prior to docking, we indicated as active residues of the A2AR the following aminoacids defining the IBS: 106,102,203,235,230,227,292,208. On the other hand, based on the analysis of multiple GPCR-G protein and GPCR-β arrestin complexes, we indicated as interacting portion of the transducers the a5 helix of mini Gs (residues: 239 to 248) and the finger loop of β arrestin 1 (residues: 63 to 76). The best binding poses were selected as those having the lowest RMSD with respect to 6TKO PDB complex.

## Acknowledgements

This work has received funding from the European Research Council (ERC) under the European Union’s Horizon 2020 research and innovation programme (“CoMMBi” ERC grant agreement No.101001784) and it was supported by a grant from the Swiss National Supercomputing Centre (CSCS) under project ID u8 and s1150. The authors also thank Dr. Francesco Saverio Di Leva from University of Naples Federico II and Dr. Stefano Raniolo from Università della Svizzera italiana (USI) for reading the paper and for the useful discussions.

## Author contributions

V.L. devised and supervised the project, V.M.D, P.C., L.M and V.L designed the simulations, V.M.D performed the calculations, developed the analytical tools and prepared the supplementary movie. All the authors analyzed the data, discussed the results and contributed to the writing of the manuscript.

## Competing interests

No competing interests are declared by the authors.

## Data and materials availability

The Path CV MetaD protocol employed in this work is available on PLUMED-NEST.

The structure of the A2AR pseudo-active state (pAs) is available as PDB (Protein Data Bank) file in the Supplementary Materials and at *www.pdbdb.com*.

The movie of the activation mechanism of A2AR in apo and ligated forms is available as Supplementary Materials and at https://www.youtube.com/watch?v=5FrtoZutSa0

## Materials & Correspondence

All correspondence and material requests should be addressed to Vittorio Limongelli.

## Notes

### Competing Interest Statement

The authors have declared no competing interest.

### Summary of Updates

Updated data discussion and revised text

