## Supplementary Information File for "Minute-timescale simulations of G Protein Coupled Receptor A2A activation mechanism reveal a receptor pseudo-active state"

### Supporting Information

#### TABLE OF CONTENTS

**Table S1:** List of atoms selected for $_{ACT}$P collective variable 2

**Table S2:** List of atoms selected for $_{TM6}$P collective variable 2

**Figure S1:** RMSD Plots of the orthosteric binding site (OBS) region of A2AR 3

**Figure S2:** RMSD Plots of the intracellular binding site (IBS) region of A2AR 4

**Figure S3:** Pearson Coefficient Matrices and PSN analysis 5

**Figure S4:** RMSD Plots of the TM helices of A2AR 6

**Figure S5:** RMSD Plots of the connector region of A2AR 7

**Figure S6:** Stability of the new pseudo-active A2AR conformation in unbiased MD 7

**Figure S7:** Plot of the IBS's volume along the *FApo* trajectory 8

**Figure S8:** Graphical representations of the Path-CVs 8

**Figure S9:** Convergence of PCV-MetaD calculations on *NECA-bound* A2AR 9

**Figure S10:** Convergence of PCV-MetaD calculations on *apo* A2AR 10

**Figure S11:** Convergence of PCV-MetaD calculations on *ZMA-bound* A2AR 11

**Figure S12:** Stability of extracted energy basins over unbiased MD simulations 12

**Figure S13:** FESs as function of $_{ACT}$P.s and $_{TM6}$P.s 13

**Figure S14:** Protein-Protein docking results 14

**Figure S15:** Projection of A2A experimental structures onto the *NECA-bound* FES 15

**Figure S15:** Comparison between $^9$F-NMR spectra and Free Energy results 15

| RMSD computation | | Alignment | |
|---|---|---|---|
| **Atom** | **Residue** | **Atom** | **Residue** |
| Cα and Cβ | 92 | Cα | 19-29 |
| Cα and Cβ | 189 | Cα | 44-64 |
| Cα and Cβ | 201 | Cα | 84-89 |
| Cα and Cβ | 205 | Cα | 119-138 |
| Cα and Cβ | 226-232 | Cα | 181-186 |
| Cα and Cβ | 234-236 | Cα | 246-251 |
| Cα and Cβ | 240-250 | Cα | 272-277 |
| Cα and Cβ | 284-289 | | |
| Cγ | 208 | | |
| Cγ | 225 | | |
| Cζ | 202 | | |
| Cζ | 288 | | |

**Table S1:** List of atoms used for the alignment and RMSD measurement of $P_{ACT}$ CV.

| Contact List | | | | | |
|---|---|---|---|---|---|
| **Contact number** | **Atom i** | **Atom j** | **$R_0$** | ***n*** | ***m*** |
| 1 | Glu228-Cδ | Arg205-Cζ | 7Å | 8 | 16 |
| 2 | Glu228-Cδ | Arg102-Cζ | 7Å | 8 | 16 |
| 3 | Glu228-Cδ | Asp101-Cγ | 16.5Å | 8 | 18 |
| 4 | Arg102-Cζ | Asp97-Cγ | 7.5Å | 6 | 18 |
| 5 | Leu235- Cγ | Leu198-Cγ | 9Å | 8 | 16 |
| 6 | Leu235- Cγ | Ile98-Cβ | 10Å | 10 | 24 |
| 7 | Tyr288-Cζ | Tyr197-Cζ | 8.5Å | 8 | 28 |
| 8 | Tyr288-Cζ | Val45-Cζ | 6.7Å | 12 | 24 |

**Table S2:** List of contacts used for computing of $P_{TM6}$ CV.

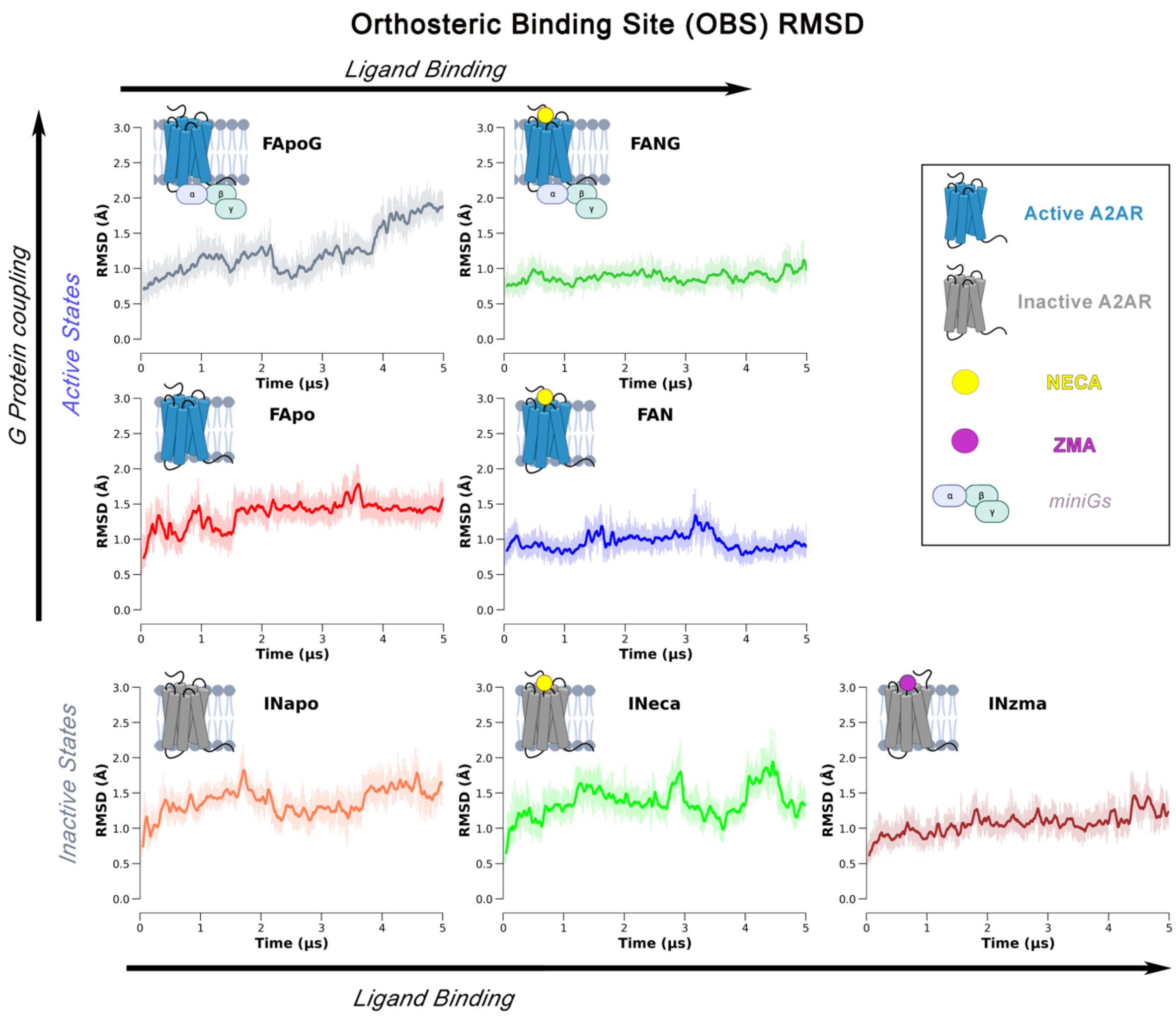


**Figure S1.** RMSD plots of the seven simulated systems in the GPCR Orthosteric binding site (OBS-Cα) with respect to the first frame of each trajectory. The bolded lines show a RMSD value smoothed with a rolling window of 5 ns, while the actual fluctuations are shown with a slight transparency.

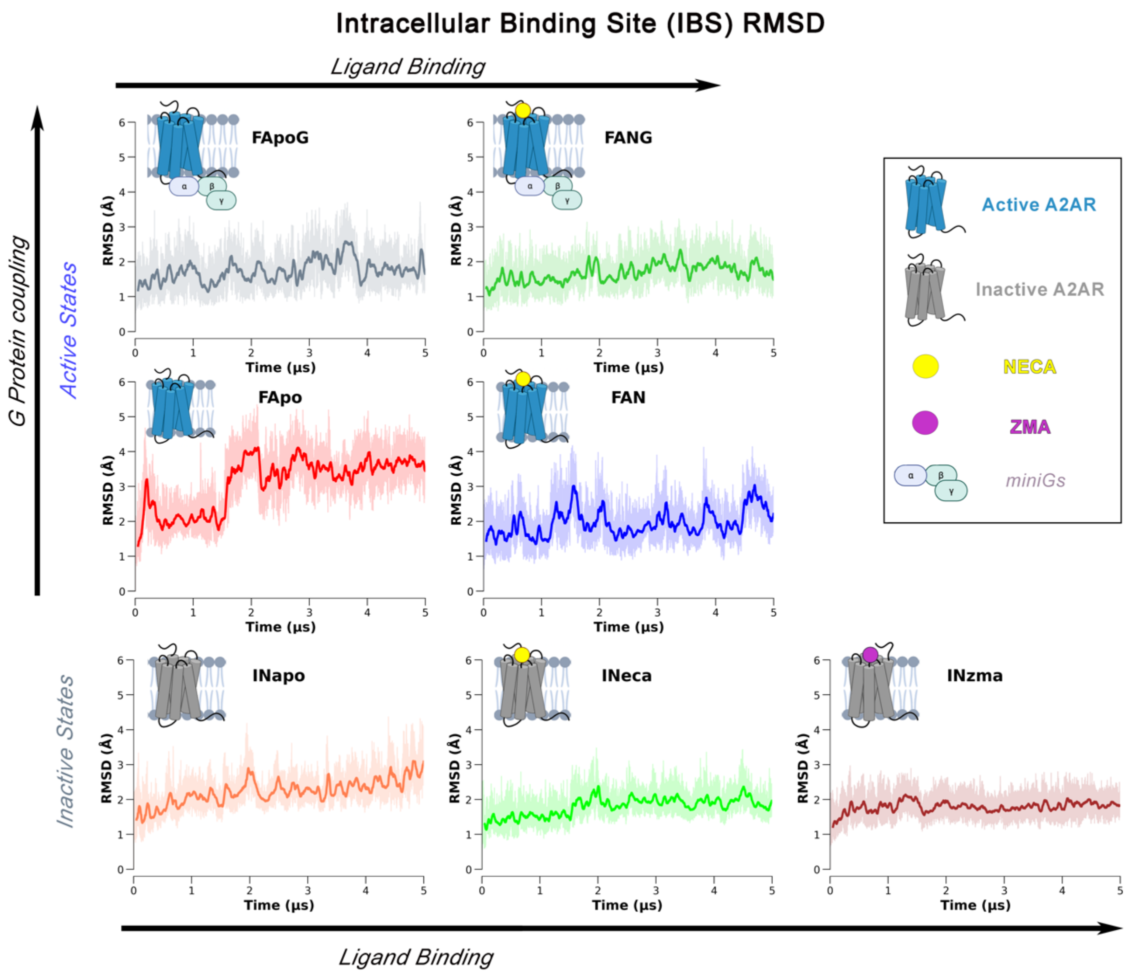


**Figure S2.** RMSD plots of the seven simulated systems in the GPCR intracellular binding site (IBS-Cα) with respect to the first frame of each trajectory. The bolded lines show a RMSD value smoothed with a rolling window of 5 ns, while the actual fluctuations are shown with a slight transparency.

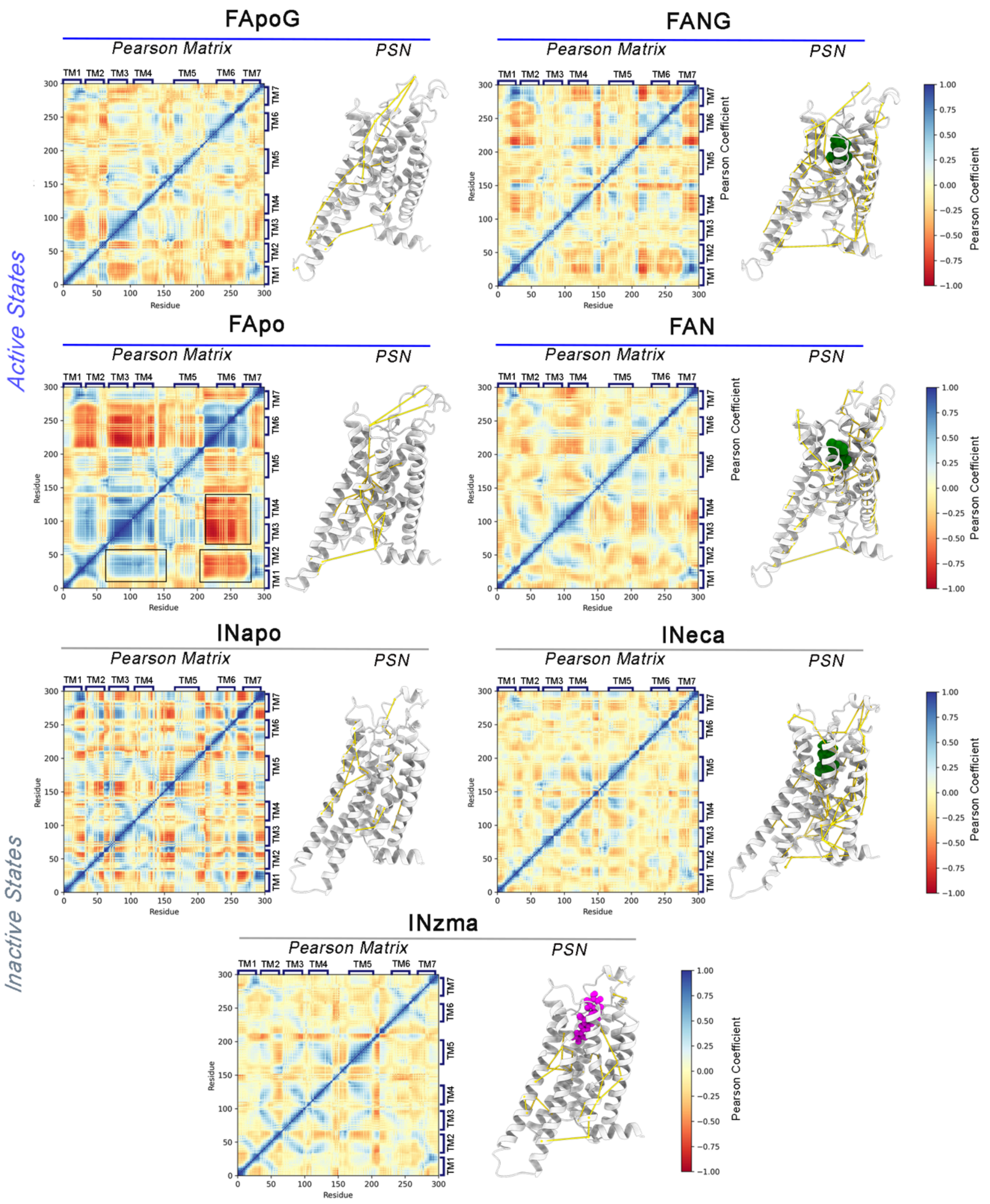


**Figure S3.** Pearson Coefficient Matrices and Protein Structure Network metapaths computed for the seven investigated MD systems. The Pearson Coefficients were computed based on the corresponding elements of the covariance matrix of the atomic (Cα of each residue) fluctuations during the unbiased trajectories. The PSN were obtained with the aid of the webpsn server; the links connecting the most important aminoacidic hubs in the global metapath are depicted as gold connections and projected onto a cartoon representation of the GPCR.

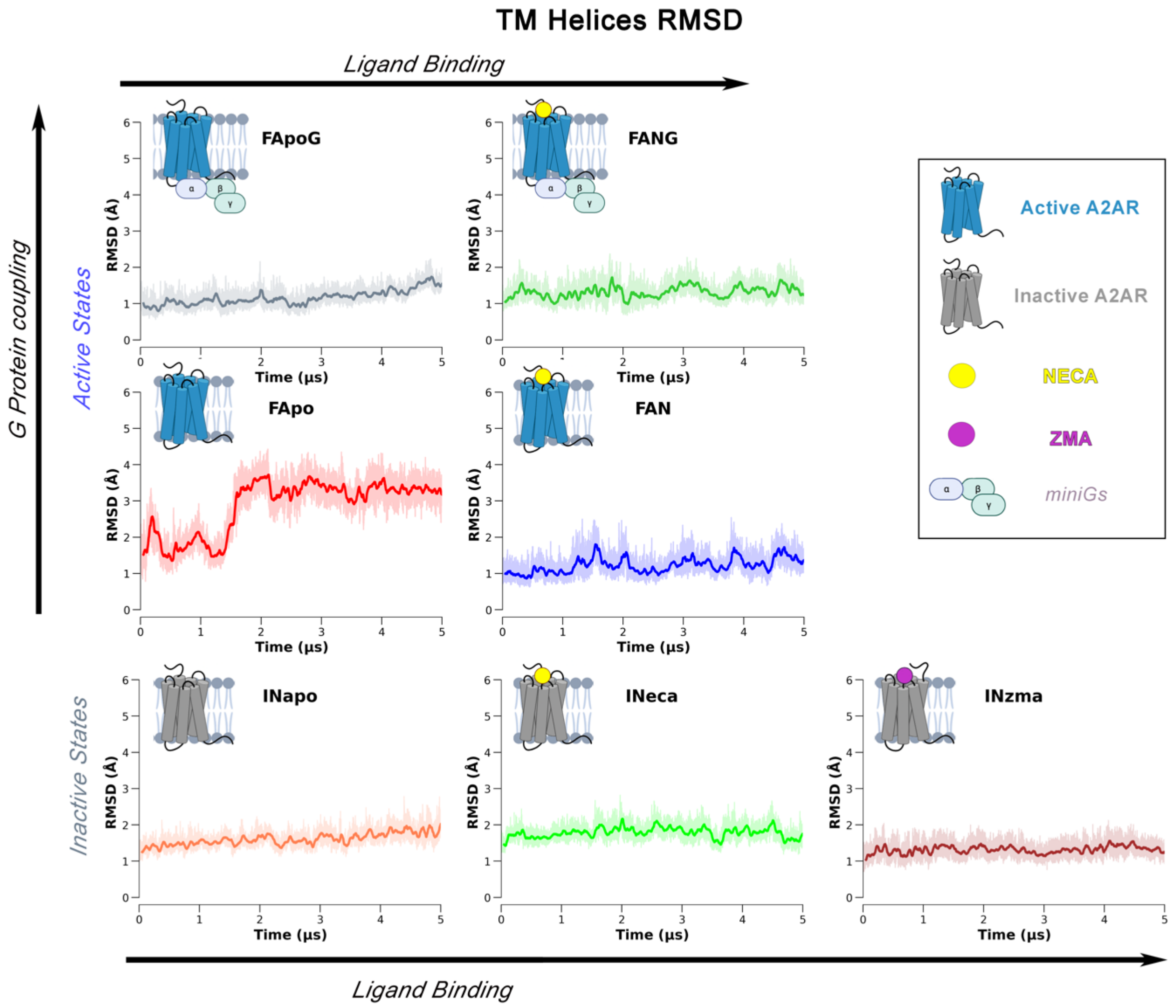


**Figure S4.** RMSD plots of the seven simulated systems computed for the GPCR transmembrane helices (Cα) with respect to the first frame of each trajectory. The bolded lines show a RMSD value smoothed with a rolling window of 5 ns, while the actual fluctuations are shown with a slight transparency.

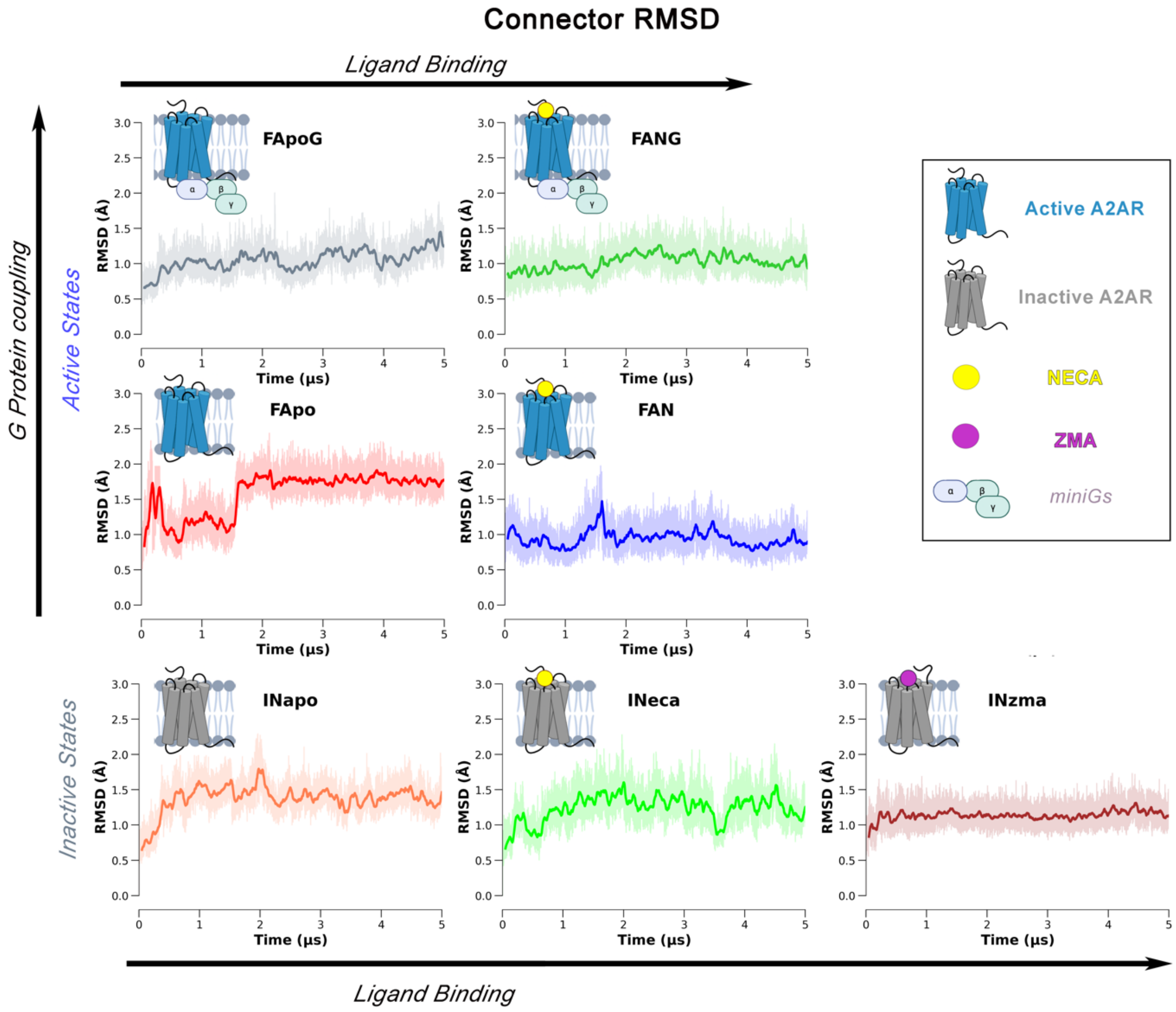


**Figure S5.** RMSD plots of the six simulated systems computed for the GPCR connector region (Cα) with respect to the first frame of each trajectory. The bolded lines show a RMSD value smoothed with a rolling window of 5 ns, while the actual fluctuations are shown with a slight transparency.

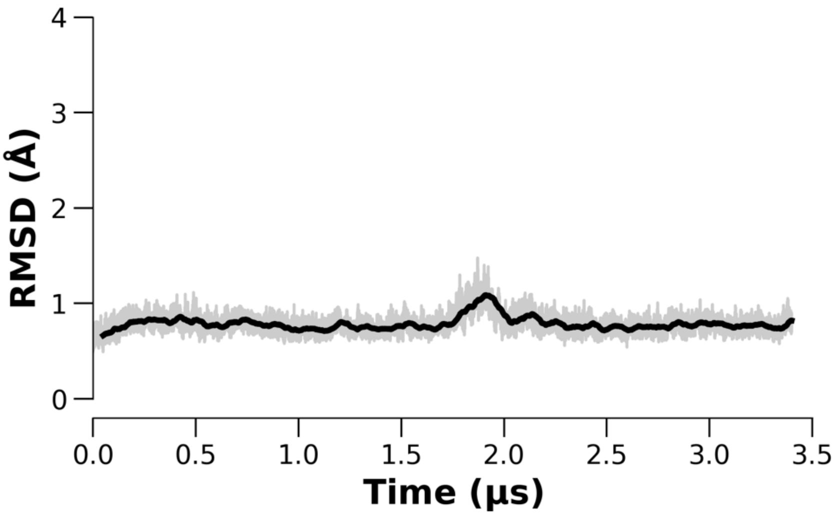


**Figure S6.** Stability of the pseudo-active A2AR in MD simulations. RMSD plots were computed for the GPCR transmembrane helices (Cα) with respect to the first frame after the conformational transition observed in the *FApo* system. The bolded lines show a RMSD value smoothed with a rolling window of 5 ns, while the actual fluctuations are shown with a slight transparency.

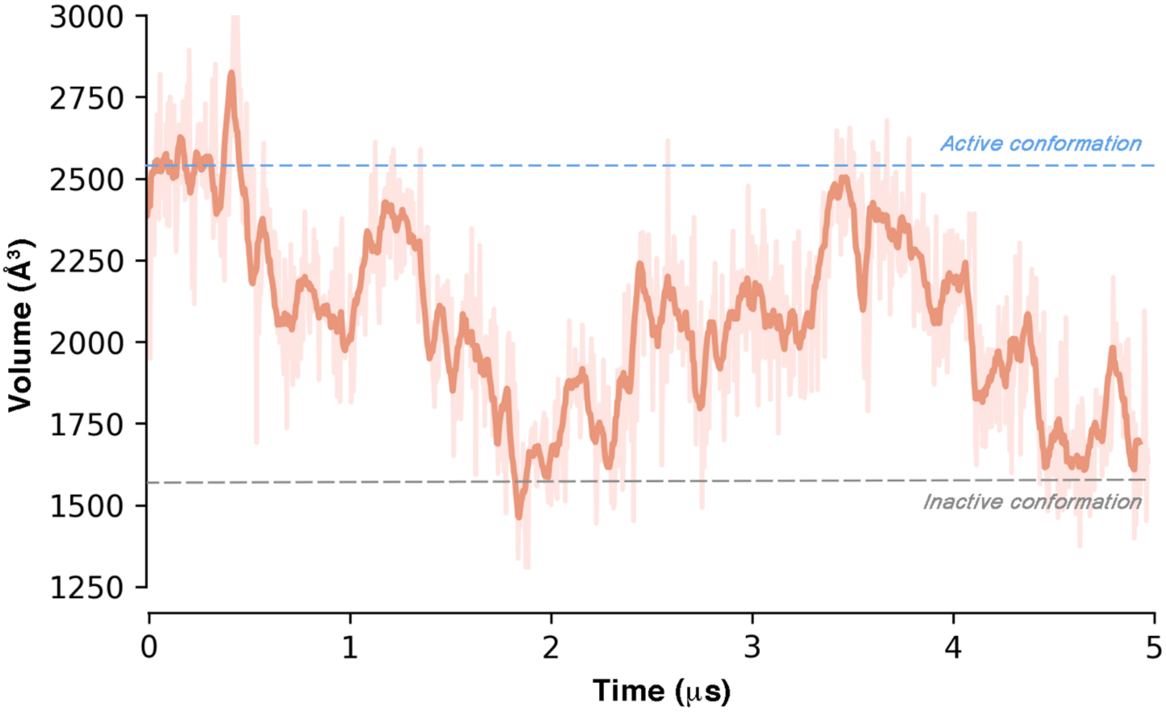


**Figure S7.** Plot of the volume computed for the IBS region along the *FApo* trajectory. The bolded lines show a volume value smoothed with a rolling window of 5ns, while the actual fluctuations are shown with a slight transparency. The reference volume values for A2AR active and inactive states are shown as light-blue and grey dashed lines, respectively. The volume computation was performed with POVME.py tool (J. Chem. Theory Comput. 2017, 13, 9, 4584–4592).

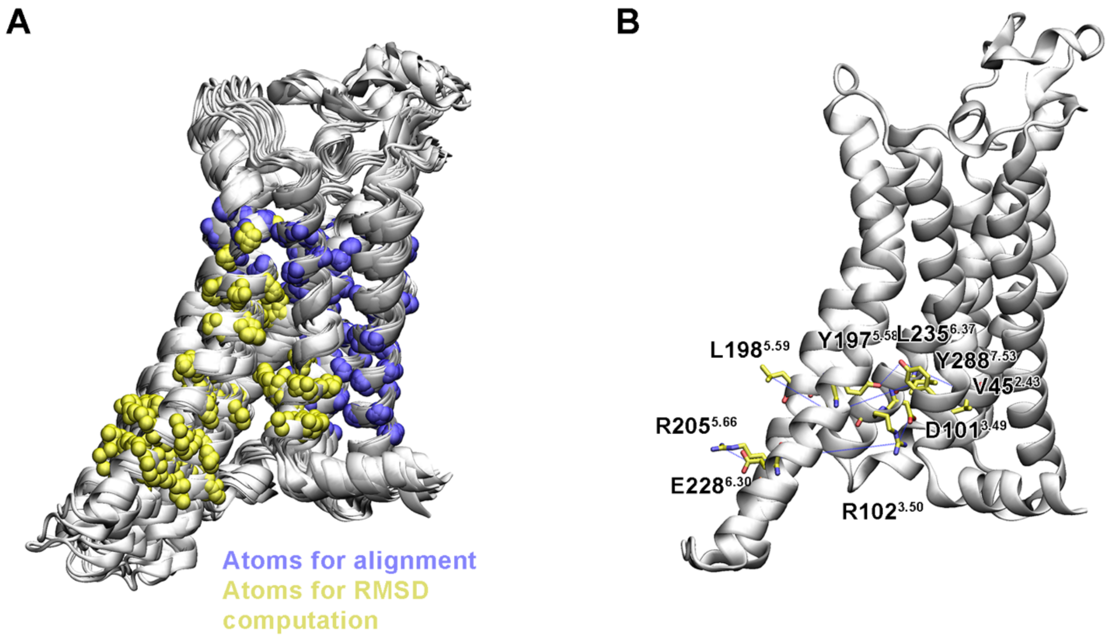


**Figure S8.** A) Cartoon representation of the 12 frames forming the reference path used for the $_{ACT}$Path Collective Variable. The RMSD matrix was computed for the atoms shown as yellow sphere and listed in Table S1, after alignment of the frames based on the position of the Cα and Cβ atoms shown as blue spheres. B) Graphical representation of the seven contacts (blue dashed lines) used for defining the $_{TM6}$Path Collective variable in the CMAP space, which are listed in Table S2. For sake of clarity the contacts are projected on the first of the nine frames forming the path, corresponding to the receptor active conformation.

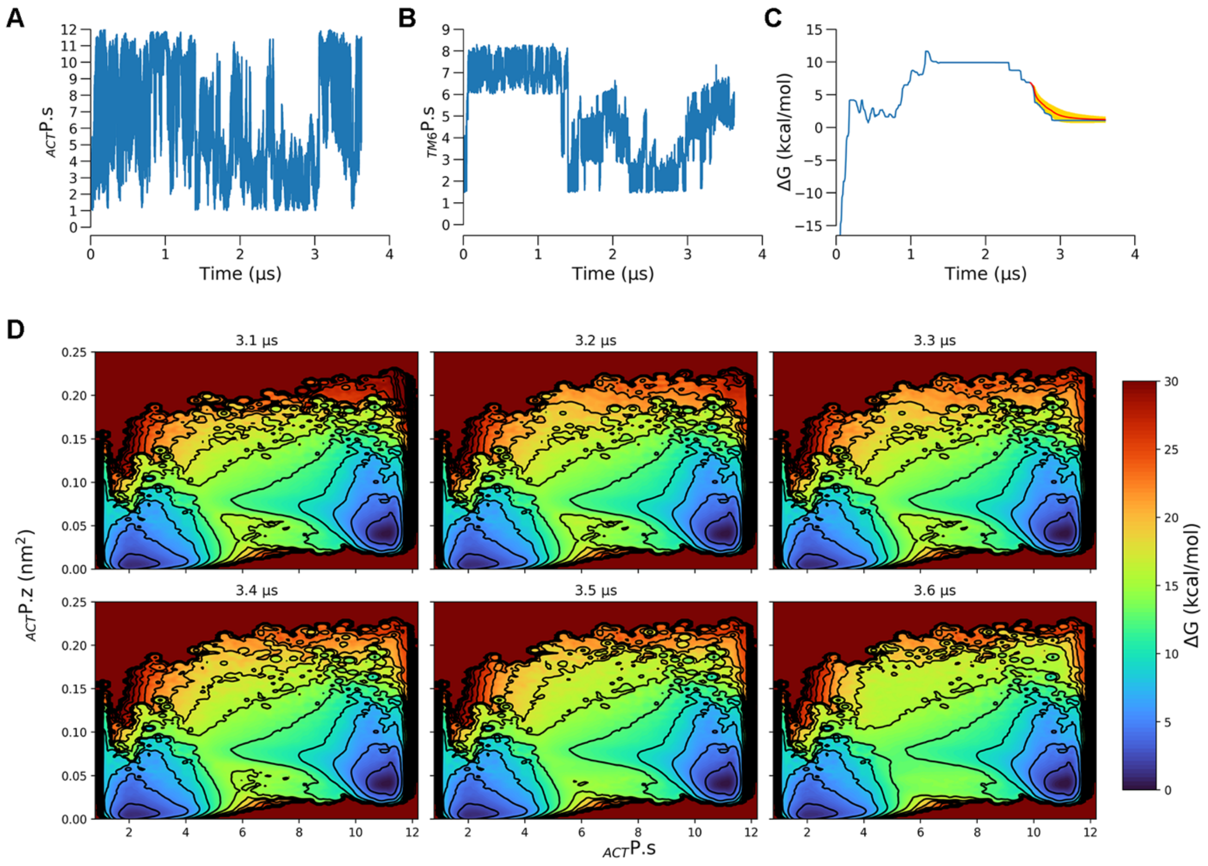


**Figure S9.** Convergence of WT-MetaD-PCVs calculation on the *NECA-bound* A2AR system. A) Time evolution of the $_{ACT}$P.s collective variable biased during the simulation. B) Time evolution of the $_{TM6}$P.s collective variable biased during the simulation. C) Free energy difference between the two main energy basins (A* and I*) as a function of the simulation time (blue). Its weighted average with the standard deviation is reported in red, error bars are shown in yellow. D) Time evolution of the reweighted FES as a function of the $_{ACT}$P.s and $_{ACT}$P.z CVs during the last 500 ns of simulation.

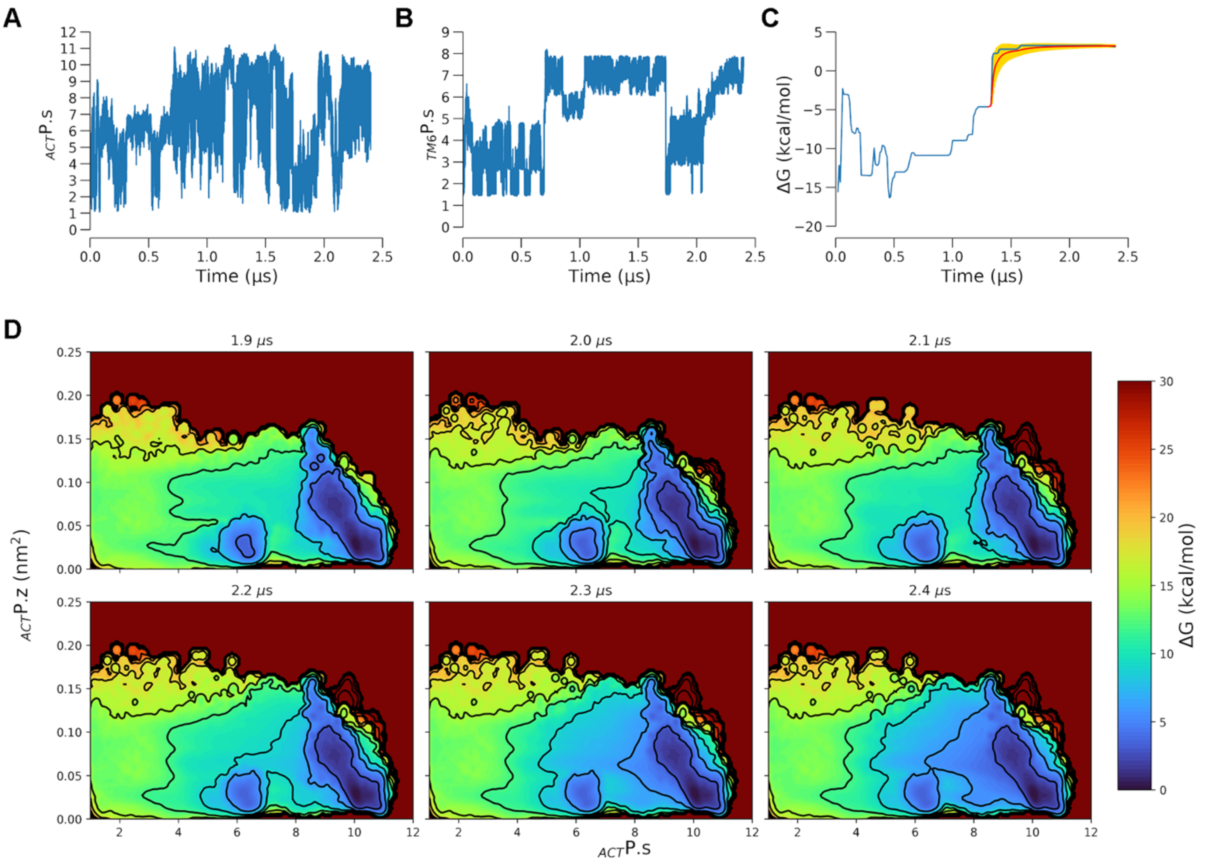


**Figure S10.** Convergence of WT-MetaD-PCVs calculation on the *apo* A2AR system. A) Time evolution of the $_{ACT}$P.s collective variable biased during the simulation. B) Time evolution of the $_{TM6}$P.s collective variable biased during the simulation. C) Free energy difference between the two main energy basins (A* and I*) as a function of the simulation time (blue). Its weighted averagesccs with the standard deviation is reported in red, error bars are shown in yellow. D) Time evolution of the reweighted FES as a function of the $_{ACT}$P.s and $_{ACT}$P.z CVs during the last 500 ns of simulation.

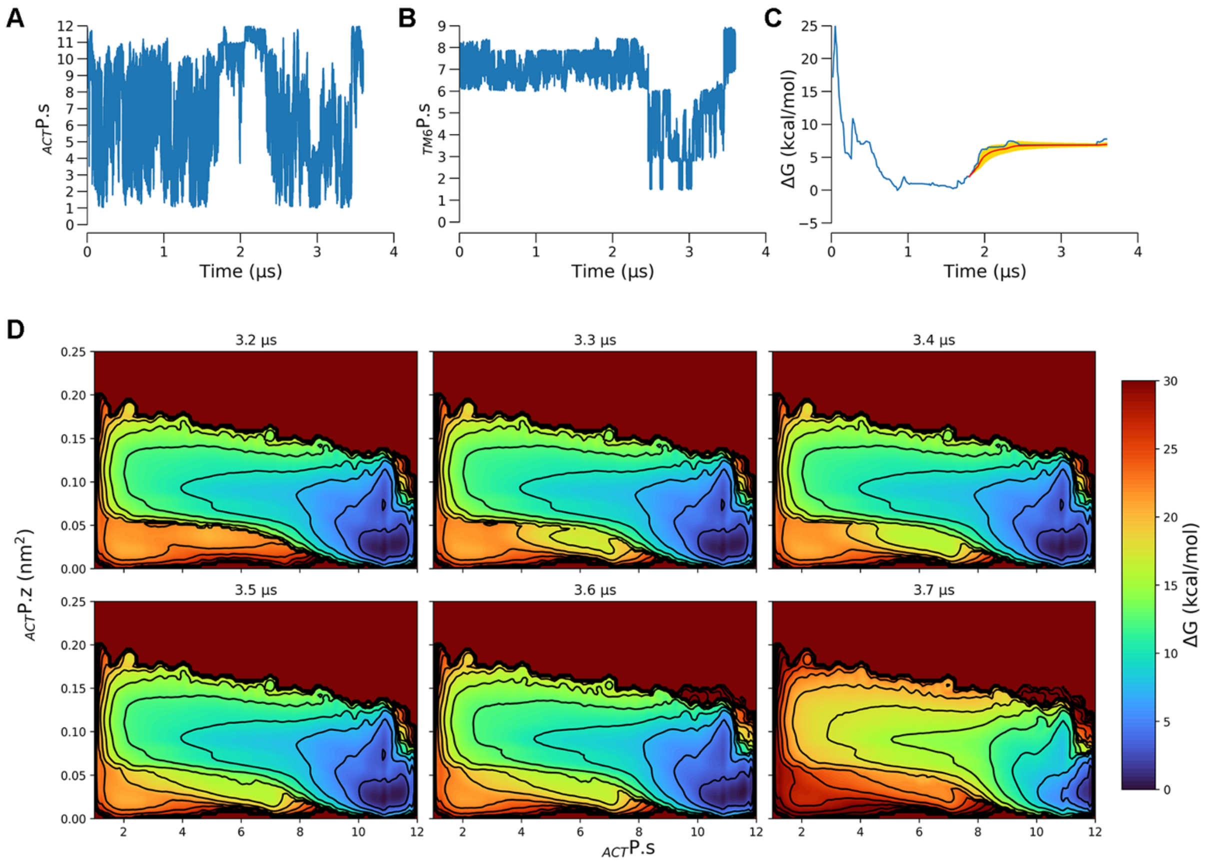


**Figure S11.** Convergence of WT-MetaD-PCVs calculation on the *ZMA-bound* A2AR system. A) Time evolution of the $_{ACT}$P.s collective variable biased during the simulation. B) Time evolution of the $_{TM6}$P.s collective variable biased during the simulation. C) Free energy difference between the two main energy basins (A* and I*) as a function of the simulation time (blue). Its weighted average with the standard deviation is reported in red, error bars are shown in yellow. D) Time evolution of the reweighted FES as a function of the $_{ACT}$P.s and $_{ACT}$P.z CVs during the last 500 ns of simulation.

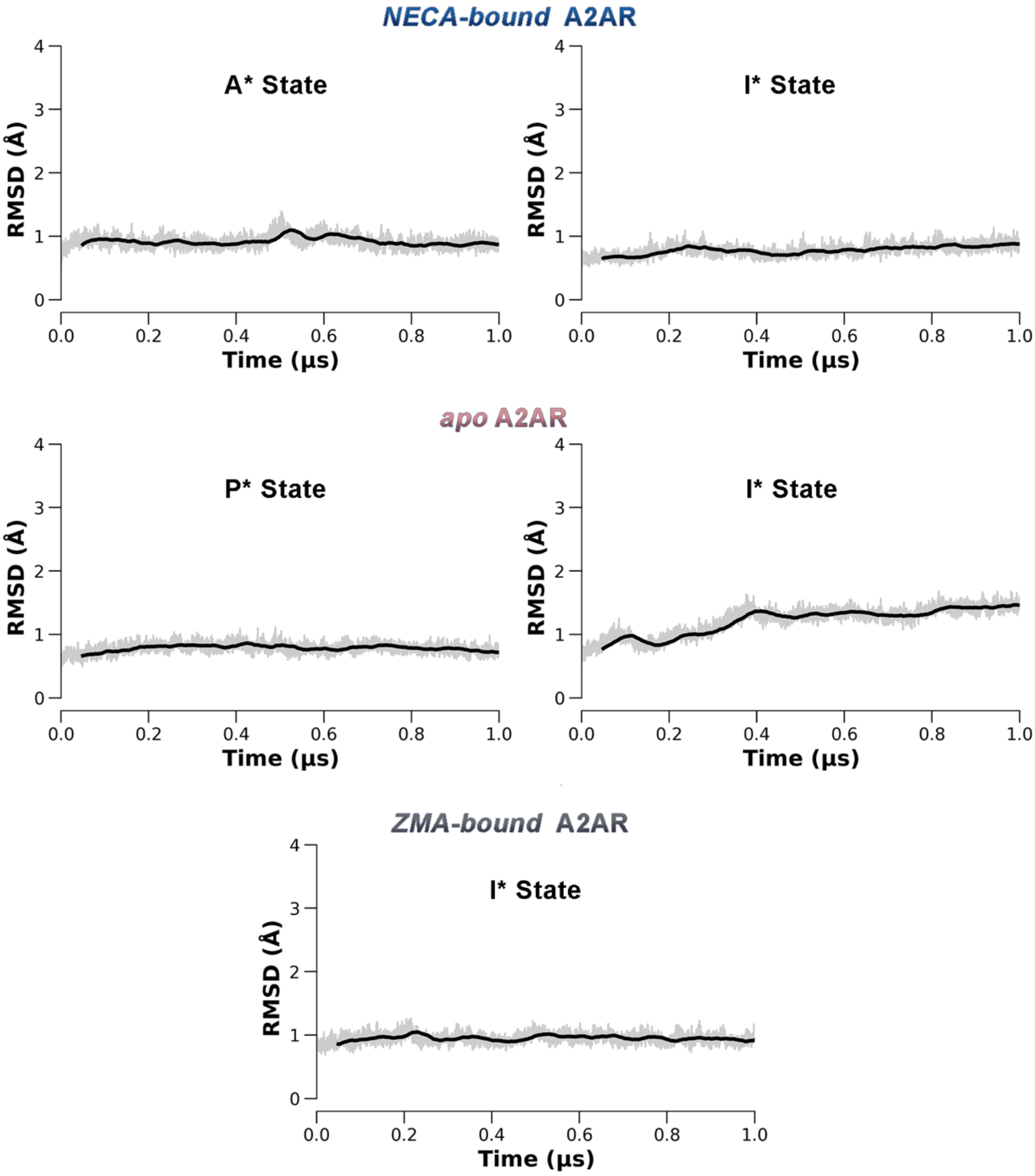


**Figure S12.** Stability of the free energy minima identified by WT-MetaD-PCVs calculations. RMSD plots are computed for the C$\alpha$ atoms of the TM helices with respect to the starting conformation of each minimum. For the A state in the *NECA-bound* system and I state in the *ZMA-bound* system, the plots refer to the first 1 μs MD simulations performed on the X-ray structures and also described in Supplementary Fig.1-4. For the P state in the *apo* system, the plot refers to the first 1 μs after the conformational transition observed in the *FApo* MD system described in Supplementary Fig. 1-4. The bolded lines show a RMSD value smoothed with a rolling window of 5 ns, while the actual fluctuations are shown with a slight transparency.

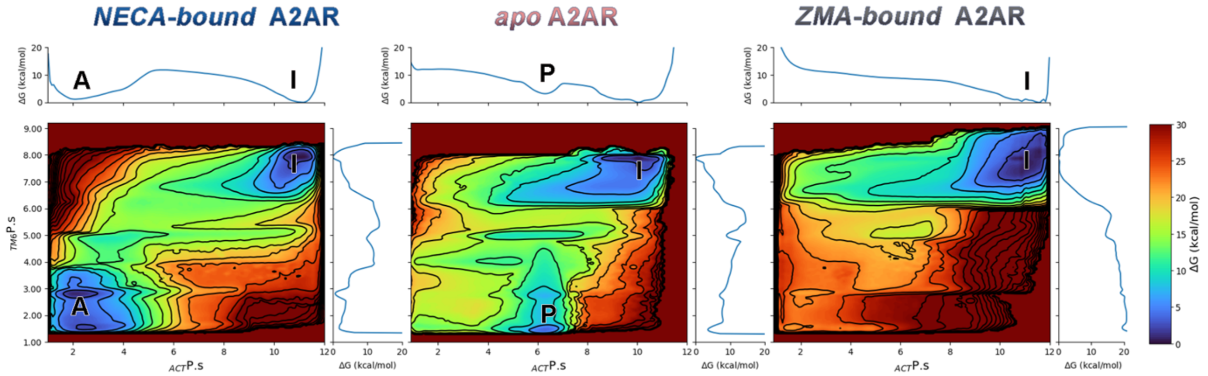


**Figure S13.** A) Activation Free Energy landscapes of the *NECA-bound*, *apo* and *ZMA-bound* forms of A2AR as a function of the $_{ACT}$P.s and $_{TM6}$P.s collective variables. Isosurfaces are displayed every 3 kcal/mol.

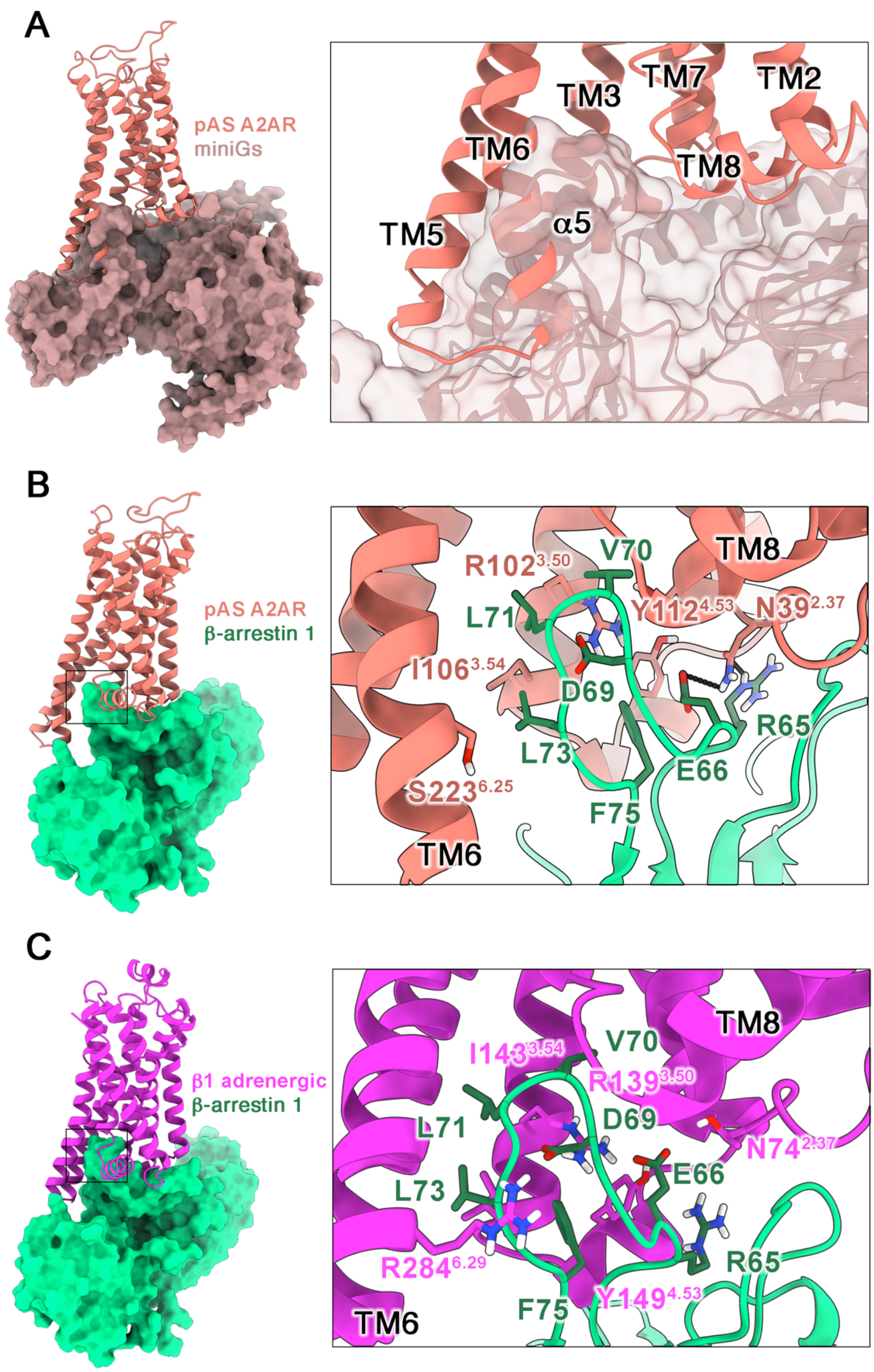


**Figure S14.** A) Steric clash between the intracellular portion of the pAS-A2AR and the α5 helix of Gα subunit. B) Docking complex predicted by Haddock of the pAS A2AR/β-arrestin 1 interaction. The pAs A2AR structure is shown as salmon cartoon, while the mini Gs and β-arrestin 1 are depicted (cartoon and surface) in brown and green, respectively. C) Experimental Cryo-EM structure of β arrestin 1 (green surface and cartoon) at the β1 adrenoreceptor IBS (purple cartoon, PDB code: 6TKO).

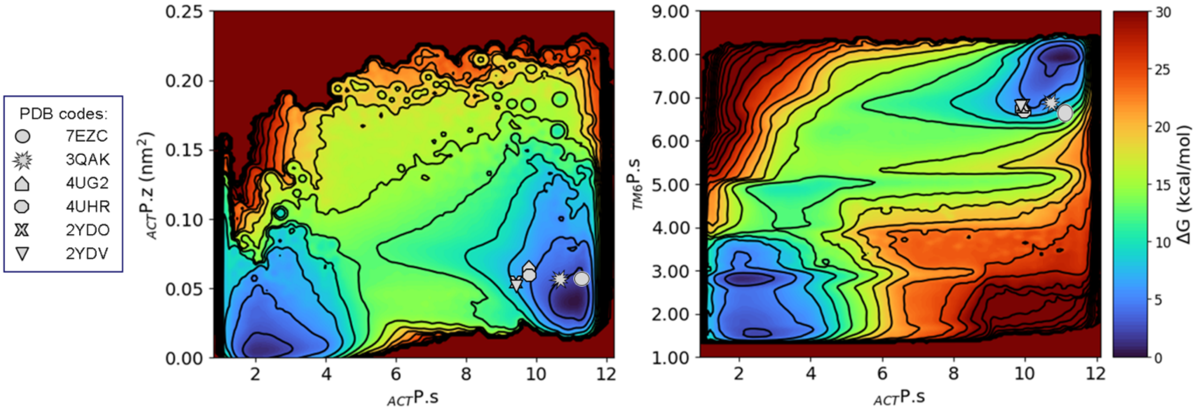


**Figure S15.** Projection of all the agonist-bound experimental structure (gray shapes) of the *uncoupled* A2AR on the activation free energy surfaces predicted in the present work for the NECA-bound system.

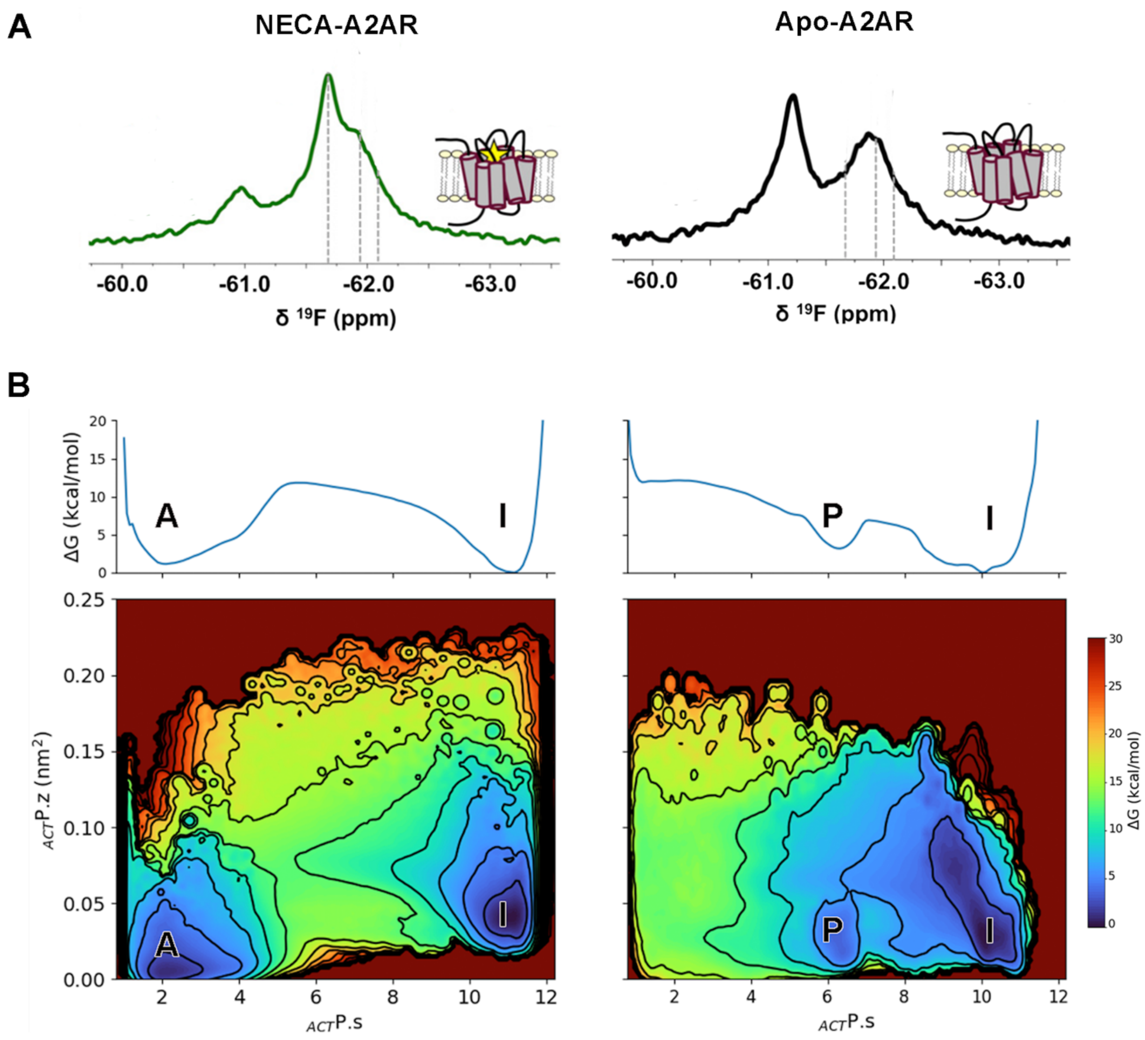


**Figure S16.** A) $^{9}$F NMR spectra of A2AR-V229C in *apo* conditions and in presence of NECA reported by Prosser and coworkers (*Cell,* **2021**, 184, 1884–1894). B) Activation Free Energy surfaces of the *NECA-bound*, *apo* and *ZMA-bound* A2AR as a function of the $_{ACT}$P.s and $_{ACT}$P.z collective variables computed by means of PCV-MetaD.
